# Mind the viscous modulus: The mechanotransductive response to the viscous nature of isoelastic matrices regulates stem cell chondrogenesis

**DOI:** 10.1101/2023.03.06.530938

**Authors:** Matthew Walker, Eonan William Pringle, Giuseppe Ciccone, Manlio Tassieri, Delphine Gourdon, Marco Cantini

## Abstract

The design of hydrogels as mimetics of tissues’ matrices typically disregards the viscous nature of native tissues and focuses only on their elastic properties. In the case of stem cell chondrogenesis, this has led to contradictory results, likely due to unreported changes of the matrices’ viscous modulus. Here, by employing isoelastic matrices with a Young’s modulus of ~12 kPa, we demonstrate that variations in viscous properties alone (i.e., loss tangent between 0.1-0.25) are sufficient to drive efficient growth factor-free chondrogenesis of human mesenchymal stem cells, both in 2D and 3D cultures. The increase of the viscous component of RGD-functionalised polyacrylamide or polyethylene glycol maleimide hydrogels promotes a phenotype with reduced adhesion, alters mechanosensitive signalling, and boosts cell-cell contacts. In turn, this upregulates the chondrogenic transcription factor SOX9 and supports neocartilage formation, demonstrating that the mechanotransductive response to the viscous nature of the matrix can be harnessed to direct cell fate.

---

The properties of tissue-mimetic hydrogels, including mechanical and biochemical cues, regulate cell adhesion and matrix secretion (1). In several recent studies, these properties have been tuned for the engineering of articular cartilage tissue, by harnessing the chondrogenic potential of mesenchymal stem cells (MSCs) (2). However, most works disregard the dynamic and dissipative nature of native extracellular matrices (ECMs) and account only for their elastic character. Biological tissues are instead viscoelastic, exhibiting time-dependent stress relaxation in response to an applied strain (3). Cells are highly sensitive to the factors governing these stress relaxation processes. Indeed, it has been demonstrated that the viscoelasticity of the substrate influences stem cell differentiation and holds a great unexplored potential in controlling stem cell chondrogenesis for cartilage engineering (4, 5). Only a few studies have addressed the effect of hydrogels’ viscous behaviour on stem cell chondrogenesis, and, in those cases, the viscous contribution of the materials was accompanied by a variation in their elastic properties or by the confounding effect of exogenous growth factors (2, 6–8). While these reports indicate that the viscous component plays a role in chondrogenesis, uncoupling the viscous contribution from other mechanical or biochemical effects is still missing in literature, and it is of crucial importance for furthering our understanding of how this role unfolds. Gong et al. suggested that the viscous character of the substrate influences cell response depending on the value of the elastic modulus of the material (9). In the case of compliant (low rigidity) materials, faster stress relaxation processes are thought to increases cell spreading, focal adhesion (FA) formation and nuclear yes-associated protein 1 (YAP) translocation to the nucleus (10), compared to slow-relaxing ones. Instead, in stiff (high rigidity) environments, fast-relaxing materials are correlated with a rapid viscous dissipation of cell-generated traction forces, reducing spreading and actin stress fibres’ organisation compared to slow-relaxing counterparts (11). Moreover, the viscous character of hydrogels has also been associated with chondrogenesis when mature chondrocytes are used for cartilage engineering instead of MSCs. Indeed, faster relaxing gels promoted secretion of an interconnected cartilage matrix by bovine articular chondrocytes; slower relaxing gels, instead, restricted this chondroinductive process (12).

MSCs are a well-established cell source for cartilage engineering as an alternative to chondrocytes. While the latter are employed in current clinical approaches to repair cartilage defects, such as autologous chondrocyte implantation (13), their use is hindered by difficulties in cell sourcing, limited proliferative capacity and possible formation of fibrocartilage (14). On the other hand, MSCs are readily expandable in culture, can be isolated from various tissue sources and have great potential in cartilage tissue engineering via their differentiation into chondrocytes (15). Indeed, MSCs are highly sensitive to the properties of their environment through integrin-mediated interactions with adhesive motifs such as Arginyl-glycyl-aspartic acid (RGD). Hence, hydrogels with optimised biomechanical and biochemical cues can direct MSC differentiation towards particular lineages (16). In terms of chondrogenesis, it has been suggested that cell spreading and strong FA attachments are not necessary or beneficial, with a low tension rounded stem cell morphology being more chondroinductive (17). An aggregated and clustered phenotype, typical of the mesenchymal condensation that occurs during cartilage development in embryogenesis, is also known to promote chondrogenesis of MSCs. This can be facilitated *in vitro* by encouraging cell-cell interactions in a highly dense 3D cellular environment (18), targeting key chondrogenic signalling events associated with N-cadherin and β-catenin (19, 20).

Based on these considerations, we have developed a set of viscoelastic hydrogel matrices that support a chondrogenic phenotype for MSCs via targeting of mechanosensitive pathways. More specifically, we have designed RGD-functionalised matrices for 2D and 3D studies using polyacrylamide (PAAm) and polyethylene glycol maleimide (PEG-MAL) hydrogels with isoelastic moduli of ~12 kPa and variable viscous component (here reported in terms of loss tangent “tan(*δ*)” varying between 0.1 and 0.25). Collectively, our results indicate that in environments with a high tan(*δ*), MSCs had a rounded phenotype with fewer FAs, lower traction forces, decreased expression of integrin β1 and β3, and hindered YAP nuclear translocation compared to an elastic (low tan(*δ*)) matrix. High instances of MSC clustering were also evident when tan(*δ*) was higher, correlating with increased N-cadherin and decreased β-catenin expression. Ultimately, early and late chondrogenesis were promoted at higher tan(*δ*) through increased expression of SOX9, collagen type II and aggrecan, and decreased expression of Runx2, fibrocartilage and chondrocyte hypertrophy markers. We believe that MSC chondrogenesis can be harnessed simply by controlling the hydrogels’ viscous component, providing an environment that facilitates a neocartilage phenotype through regulation of cell adhesion, mechanotransduction and cell-cell communication.

## Results

### Fabrication and characterisation of isoelastic hydrogels with variable viscous component

PAAm hydrogels are widely used for 2D cell studies due to their highly tuneable mechanical properties and ease of functionalisation with ECM peptides, such as RGD, to promote cell adhesion (21). They can be fabricated with controlled viscoelastic properties using various strategies, such as incorporating linear, high molecular weight PAAm chains (22). Here, we optimised an alternative strategy to tune PAAm viscoelasticity by fabricating gels with high polymer content and relatively low crosslinking, encouraging physical entanglement of polymer chains (**Figure 1A**), as suggested previously by Cameron et al. (23). PEG-MAL hydrogels are instead suitable for 3D cell studies due to their lack of toxic precursor components, which are present in PAAm hydrogels (24). The maleimide groups of PEG-MAL are crosslinked and functionalised with thiolated peptides at physiological pH through Michael-type addition, forming bioactive and cytocompatible hydrogels (25). We used 8-armed PEG-MAL to fabricate hydrogels and controlled their viscoelastic properties by adjusting polymer molecular weight (**Figure 1A**); this strategy has also been effective with other material systems such as alginate (26). The specific compositions of each hydrogel can be found in the **Materials and Methods** section.

**Figure 1.**
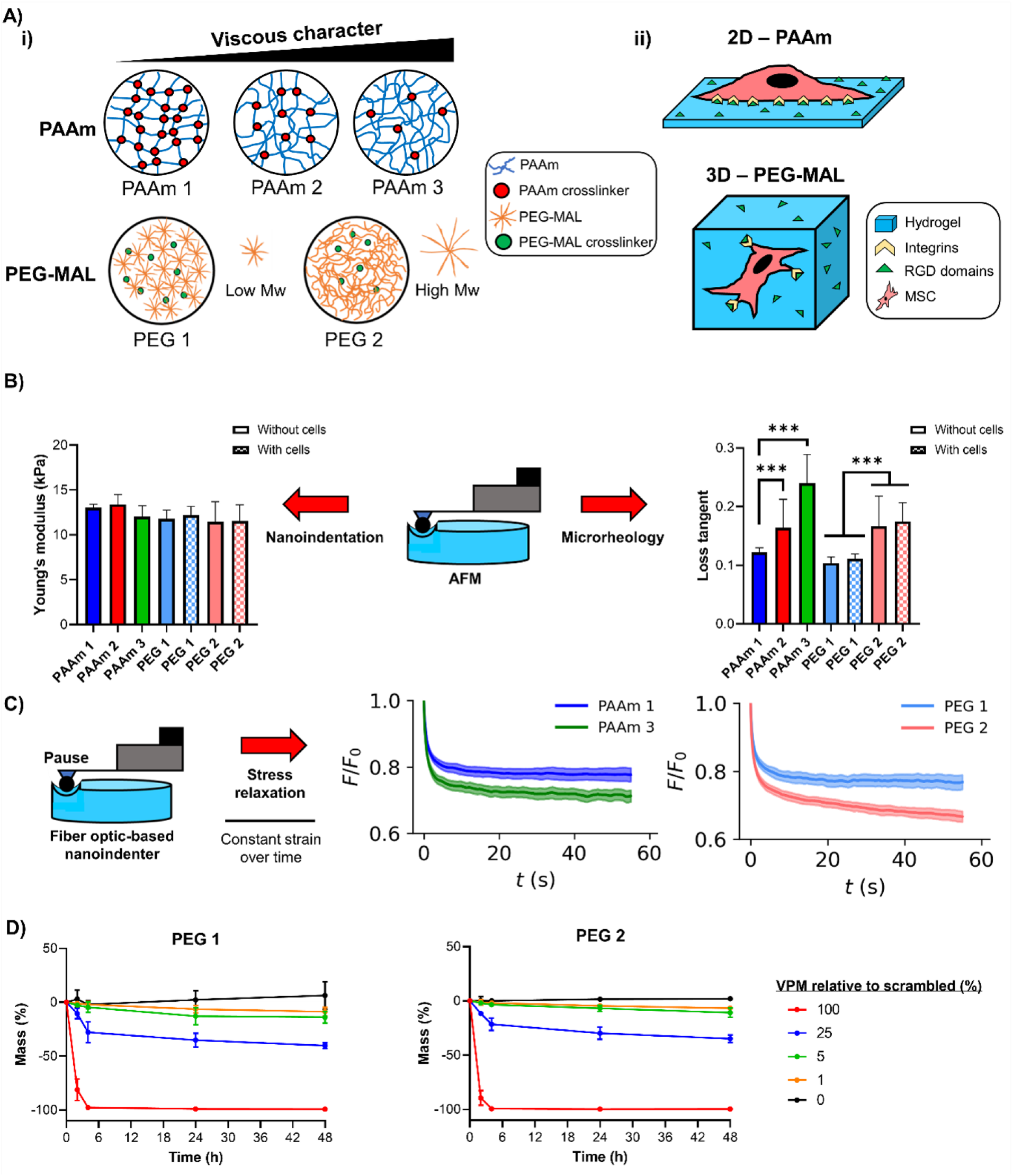
PAAm and PEG-MAL isoelastic hydrogels with tuneable viscous component. **A)** Representation of PAAm and PEG-MAL polymer:crosslinking ratios and molecular weight differences within hydrogels and influence on their loss tangent (**i**) and sketch of 2D/3D dimensionality of each hydrogel system as cell culture platforms (**ii**). **B)** AFM microscale measurements of PAAm and PEG-MAL (crosslinked with 1% VPM relative to scrambled) hydrogels using nanoindentation for Young’s modulus (left) and microrheology for loss tangent (right); PEG-MAL hydrogels were measured without cells and in the presence of encapsulated cells; n=50-100. **C)** Stress relaxation measurements of hydrogels using a fibre-optic based nanoindenter with pause step and quantification of stress relaxation curves for PAAm (left) and PEG-MAL (crosslinked with 1% VPM relative to scrambled) (right) hydrogels, n=100. **D)** Biodegradability of PEG-MAL hydrogels crosslinked with different degrees of VPM (relative to scrambled VPM) represented by mass loss over 48 hours in the presence of collagenase D for PEG 1 (left) and PEG 2 (right), n=3. For all figures, data are represented as mean ± standard deviation and differences are considered significant for p≤ 0.05 using one-way ANOVAs for multiple comparisons (*** p ≤ 0.001).

By using atomic force microscopy (AFM), we found that all PAAm and PEG-MAL hydrogels had similar Young’s moduli of ~12 (± 1) kPa through nanoindentation measurements. Additional microrheology testing revealed loss tangent values ranging between ~0.1 and ~0.25 (**Figure 1B**). We also observed that the encapsulation of MSCs within PEG-MAL hydrogels did not significantly impact the gels’ mechanics (**Figure 1B**). Bulk rheology measurements confirmed that the hydrogels displayed similar shear elastic moduli and variable viscous moduli, as shown in **Supplementary Figure 1A**. The viscoelastic properties of the hydrogels were further assessed by stress relaxation measurements, which revealed that PAAm and PEG-MAL hydrogels with higher loss tangent values displayed faster stress relaxation (**Figure 1C**) and higher energy dissipation (**Supplementary Figure 1B**), in agreement with previous studies (10, 12, 26). We were also able to control the biodegradability of the PEG-MAL hydrogels by adjusting the ratio of the protease-sensitive peptide GCRDVPMSMRGGDRCG (VPM) relative to a scrambled non-degradable counterpart during crosslinking of the hydrogels (**Figure 1D**). We observed that adjusting hydrogel degradability did not influence swelling behaviour, indicating that the macromolecular structure and porous network of the hydrogels was unaffected (**Supplementary Figure 2A, B**). Finally, AFM imaging of all hydrogels revealed a similar topography independently of gel composition and mechanical properties, as indicated by similar roughness values (**Supplementary Figure 2C**).

### The viscous character of isoelastic matrices regulates hMSC adhesion and spreading

Hydrogels were functionalised with 2 mM RGD peptide to facilitate cell adhesion, as neither PAAm nor PEG-MAL contain naturally occurring cell binding sites. Using fluorescently labelled RGD, we observed uniform ligand densities across the PAAm hydrogels’ surfaces, indicating similar availability of cell adhesion sites (**Supplementary Figure 3**). We also verified that cells were viable when encapsulated within the PEG-MAL hydrogels, confirming their suitability for 3D culture (**Supplementary Figure 4**). Given the aim of this paper, and the successful achievement of isoelastic hydrogels, from now on all samples are discriminated by means of their relative loss tangent tan(*δ*) values; moreover, for PEG-MAL hydrogels, 1% of the degradable crosslinker VPM is used unless stated otherwise.

After seeding human MSCs (hMSCs) on the surface of PAAm hydrogels, we observed striking differences in their spreading behaviour and FAs size and number. On substrates with a higher tan(*δ*) cell spreading was diminished, as demonstrated by reduced cell area and increased circularity (**Figure 2Ai, ii**); this coincided with a reduction in phosphorylated-focal adhesion kinase (p-FAK) intensity, average FA length, and frequency of FAs over 2 μm (**Figure 2Aiii**). Previous work has shown that a rounded MSC phenotype encourages chondrogenesis in 2D with low, non-localised vinculin expression (17). We observed the same behaviour in cell spreading and circularity for hMSCs on 2D PEG-MAL hydrogels with increased tan(*δ*) (**Supplementary Figures 5 and 6C**). Analysis of various hMSC cytoskeletal shape descriptors over time showed results consistent with our observations of cell spreading behaviour (**Supplementary Figure 6**). These observations agree with previous works reporting that cells display reduced spreading on stiffer hydrogels as tan(*δ*) increases (9–11). It has been suggested that this behaviour could be related to rapid energy dissipation of cell-generated traction forces into the matrix, which hinders spreading and activation of mechanoresponsive signalling pathways (11). Indeed, we observed decreased actin fibres anisotropy for hMSCs on more viscous PAAm gels, implying a reduced cytoskeletal organisation of cells on those substrates (**Supplementary Figure 6B**). Correspondingly, hMSCs applied significantly lower traction forces on PAAm hydrogels with a higher loss tangent value than on more elastic ones (**Figure 2B**). Moreover, the expressions of integrins β1 and β3 (known RGD receptors) were significantly reduced for hMSCs interacting with higher tan(*δ*) hydrogels, indicating less integrin availability for cell attachment via RGD-integrin interactions, both on 2D PAAm hydrogels and within 3D PEG-MAL matrices (**Figure 2C**).

**Figure 2.**
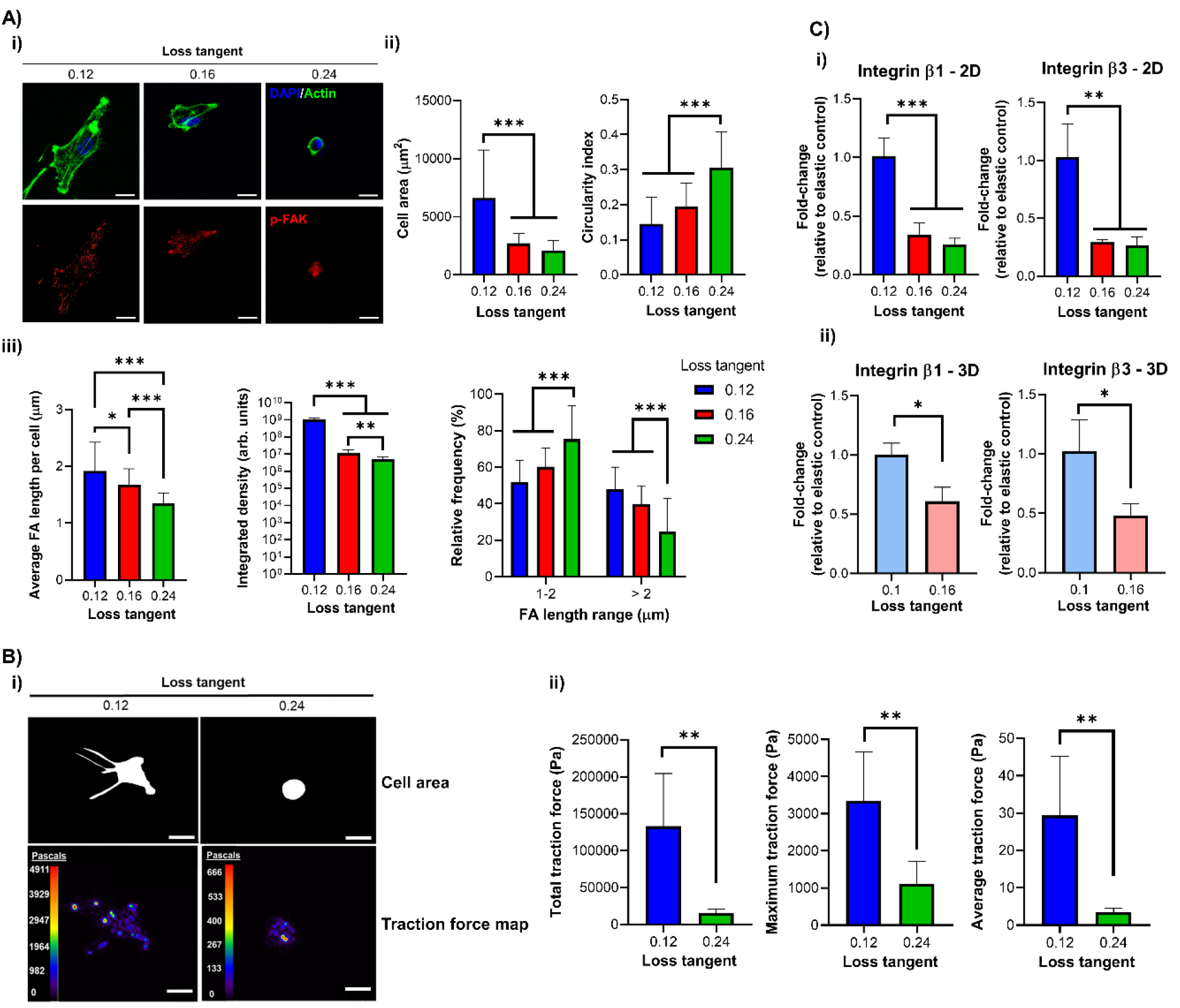
hMSC adhesion and spreading decrease as the matrix viscous component increases. **A)** Representative immunofluorescence images of hMSCs cultured for 24 hours on 2D PAAm hydrogels with DAPI (blue), actin (green) and p-FAK (red) staining (**i**) with quantification of cell area (left) and circularity (right), n=31-35 (**ii**), and quantification of average FA length per cell (left), p-FAK signal intensity (middle) and relative frequencies of FAs between 1-2 μm and > 2 μm (right), n=25-50 (**iii**). **B)** Representative images of the cell mask area and traction stress maps with cell traction stresses (in Pascals) for hMSCs cultured for 24 hours on 2D PAAm hydrogels (**i**) and quantification of the total (left), averaged (middle) and maximum (right) traction stresses, n=5 (**ii**). **C)** qPCR data from hMSCs cultured on 2D PAAm (**i**) and in 3D PEG-MAL gels (**ii**) for 3 days showing fold-change in gene expression of integrin β1 (left) and β3 (right) relative to control with stronger elastic character (0.12/0.1 loss tangent samples) and normalised to GAPDH, n=3. For all figures, data are represented as mean ± standard deviation and differences are considered significant for p≤ 0.05 using one-way ANOVA or t-tests for multiple or pairwise comparisons respectively (* p ≤ 0.05, ** p ≤ 0.01, *** p ≤ 0.001). All hydrogels were functionalised with 2 mM RGD peptide to allow cell adhesion. All PEG-MAL hydrogels were crosslinked using peptide ratios of 1% VPM and 99% scrambled VPM. Scale bars = 20 μm.

### hMSC mechanotransduction is regulated by the viscous component of isoelastic matrices

Next, we investigated whether the viscous component of our matrices also affected mechanotransduction, driving transcriptional control through regulators such as YAP (4). YAP has been identified in previous work as a negative regulator of chondrogenesis in MSCs (27); however, the role of mechanosensitive YAP signalling is relatively unexplored in hydrogel-driven chondrogenesis, despite its established role as a mechanical rheostat (28). Here, we observed decreased nuclear YAP localisation on PAAm hydrogels with a higher loss tangent, implying reduced YAP-mediated transcriptional regulation of anti-chondrogenic target genes (**Figure 3A**). We observed the same effect for PEG-MAL hydrogels with a higher tan(*δ*), both in 2D (**Supplementary Figure 7**) and in 3D (**Figure 3B**).

ROCK and Ras-related C3 botulinum toxin substrate 1 (Rac1) signalling are important events in the regulation of cytoskeletal organisation. Inhibition of ROCK signalling via Y-27632 has been shown to cause increased chondrogenesis (7, 29); this suggests that reduced cytoskeletal tension is beneficial to facilitate a chondrogenic phenotype. Here, we observed that inhibition of ROCK or Rac1 signalling reduced nuclear YAP translocation in hMSCs on PAAm hydrogels with a stronger elastic character; combined ROCK and Rac1 inhibition reduced nuclear YAP further and facilitated a phenotype similar to that of cells on more viscous hydrogels (**Supplementary Figure 8**). On the other hand, inhibition of ROCK and Rac1 signalling had no influence on nuclear YAP in hMSCs seeded on PAAm hydrogels with a higher loss tangent; only the inhibition of both caused a slight reduction in translocation (**Supplementary Figure 8**). Collectively, these results suggest that ROCK and Rac1 signalling are significantly less active in cells on hydrogels with a high tan(*δ*), indicating a more chondrogenic environment.

Lamins are major structural and mechanotransductive proteins of the nucleus. It has been shown that MSCs on soft matrices exhibit a less spread nucleus and low levels of lamin A/C expression due to its rapid phosphorylation in response to reduced cytoskeletal tension (30). We observed that on hydrogels with a higher tan(*δ*), the ratio of lamin A/C:B1 was significantly reduced, indicating that hMSCs have a similar lamin A/C profile to that seen on soft matrices (**Figure 3C**). Additionally, we observed that the nuclei of hMSCs on hydrogels with a higher tan(*δ*) had a smaller area and lower solidity, which indicate reduced nuclear spreading and correlate with a phenotype for reduced lamin A/C expression (**Supplementary Figure 6H-M**).

**Figure 3.**
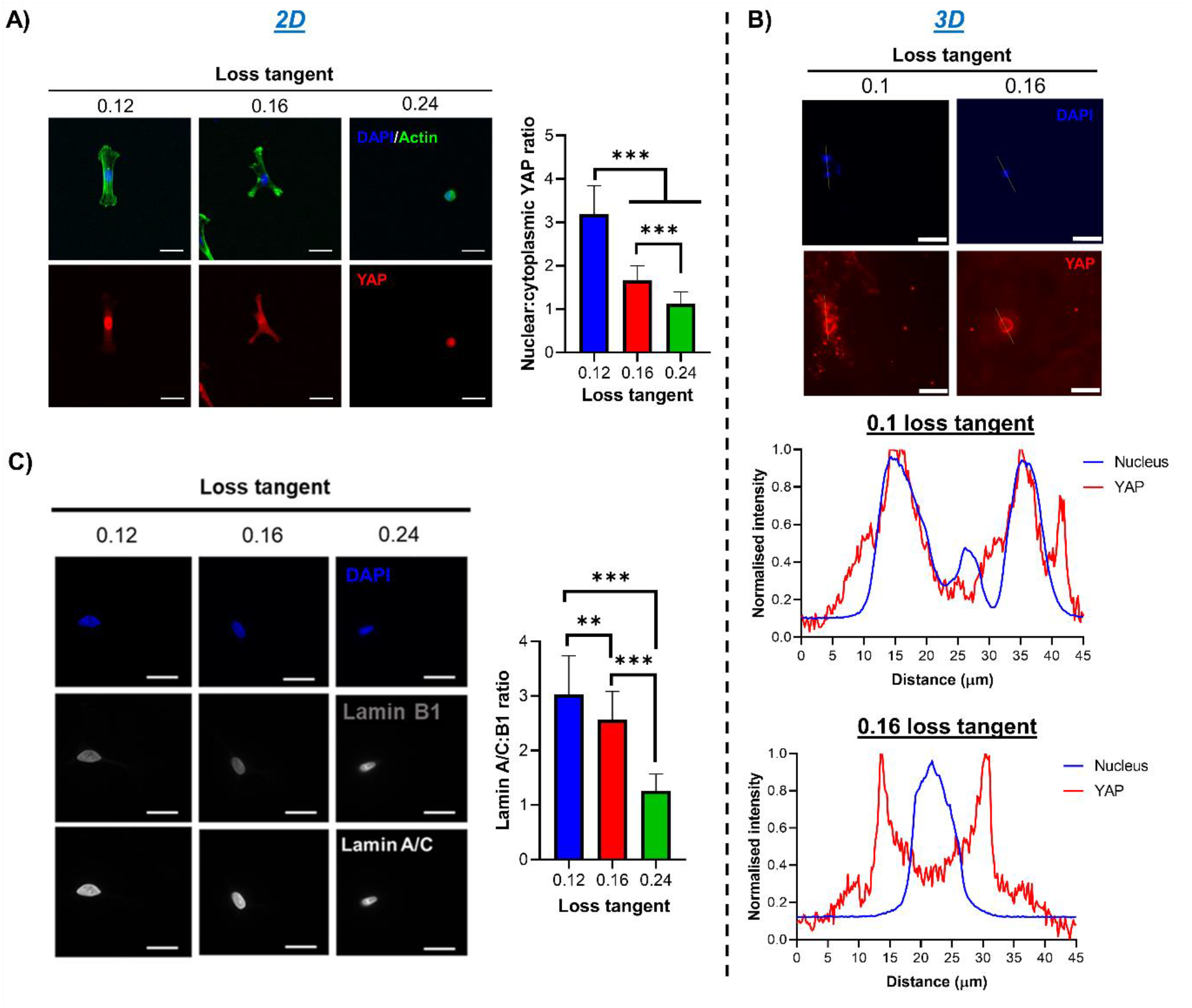
hMSC mechanotransduction is regulated by the matrix viscous component. **A)** hMSCs cultured for 3 days on 2D PAAm gels before staining for DAPI (blue), actin (green), and YAP (red), with representative immunofluorescence images (left) and quantification of nuclear:cytoplasmic YAP ratio (right), n=22-30. **B)** hMSCs cultured for 3 days in 3D PEG-MAL gels before staining for DAPI (blue) and YAP (red), with representative immunofluorescence images (top) and representative line scan analysis of nuclear and YAP intensity using ‘plot profile’ in ImageJ (bottom), showing that YAP is mainly nuclear in elastic gels and cytoplasmatic in viscoelastic ones. **C)** Representative immunofluorescence images of hMSCs cultured for 3 days on 2D PAAm hydrogels with DAPI (blue), lamin B1 and A/C staining (left) and quantification of lamin A/C:B1 ratio (right), n=30-31. For all figures, data are represented as mean ± standard deviation and differences are considered significant for p≤ 0.05 using one-way ANOVAs for multiple comparisons (** p ≤ 0.01, *** p ≤ 0.001). All hydrogels were functionalised with 2 mM RGD peptide to allow cell adhesion. All PEG-MAL hydrogels were crosslinked using peptide ratios of 1% VPM and 99% scrambled VPM. Scale bars = 20 μm.

### Cell-cell communication in hMSCs is regulated by the matrices’ viscous component

Cell-cell communication is crucial for chondrogenesis (2). Indeed, micromass/pellet cultures are used as scaffold-free chondrogenic systems by facilitating high density cell-cell contacts and mesenchymal condensation (31). Wnt signalling, involving N-cadherin and β-catenin, is a crucial event in cell-cell-mediated chondrogenesis, with canonical Wnt activation and β-catenin accumulation having been implicated as negative regulators in this process (32). Repressed Wnt/β-catenin signalling is likely to be involved in early chondrogenesis, as inhibition for 3 days was reported to increase chondrogenic gene expression (20). Here, we observed that, on PAAm gels with a higher tan(*δ*), hMSCs cultured for 3 days expressed higher levels of N-cadherin, which was localised to the cell-cell junctions within clusters of aggregated cells; minimal clustering and lower N-cadherin levels were instead observed on gels with a lower tan(*δ*) (**Figure 4A**). We also observed an increase in N-cadherin expression by qPCR as well as a concomitant downregulation in β-catenin, which is likely to facilitate early chondrogenesis through N-cadherin-mediated inhibition of β-catenin (**Figure 4B**) (19). This increased cell clustering was also seen in a 3D PEG-MAL environment with a higher tan(*δ*) (**Figure 4C**) and coincided with increased N-cadherin expression and reduced β-catenin (**Figure 4D**). It is likely that increased clustering due to viscoelastic mechanoregulation of hMSCs facilitates N-cadherin mediated inhibition of β-catenin.

**Figure 4.**
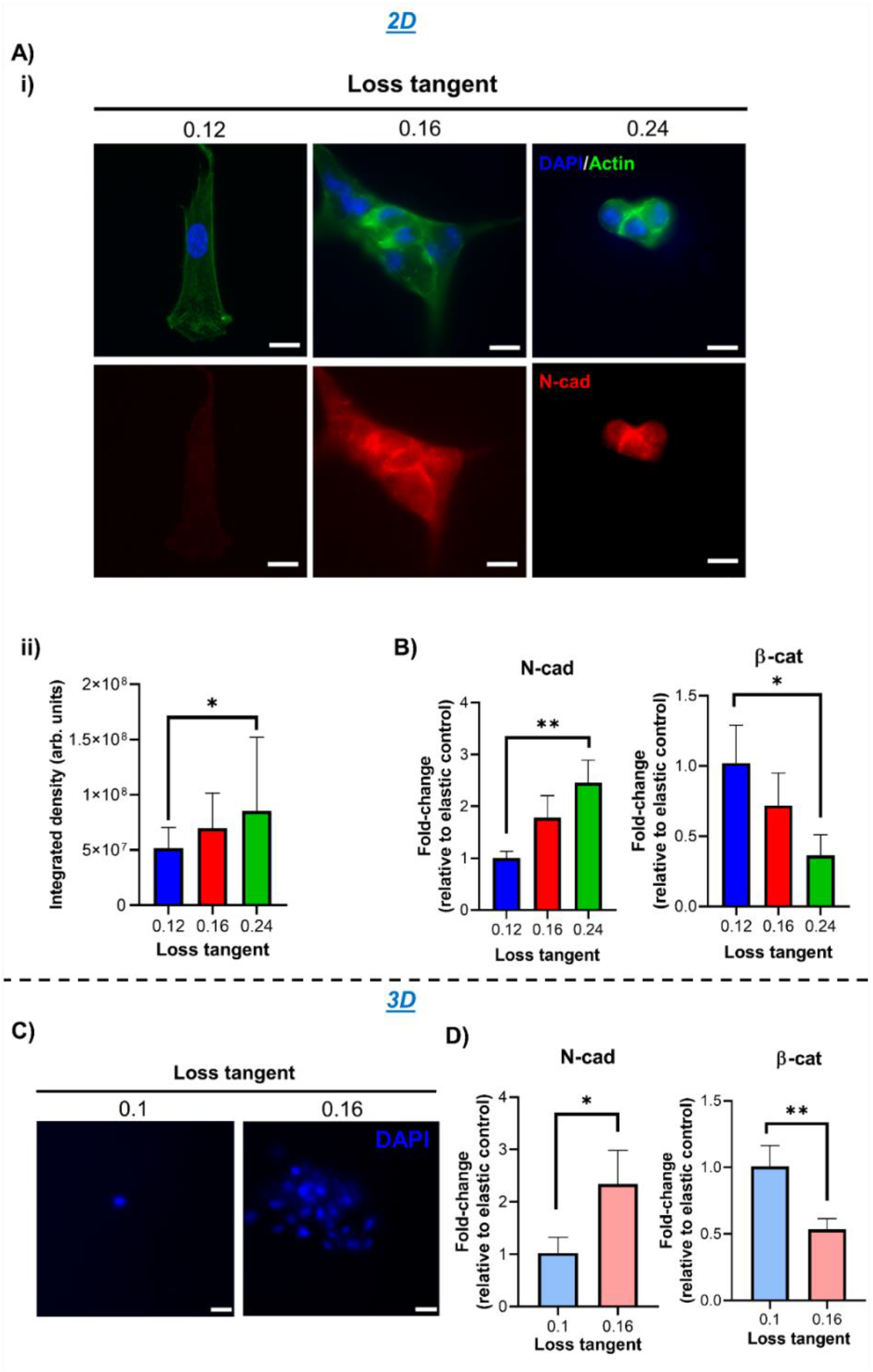
hMSC cell-cell signalling is enhanced in matrices with a higher loss tangent. **A)** Representative immunofluorescence images of hMSCs cultured for 3 days on 2D PAAm hydrogels with DAPI (blue), actin (green) and N-cadherin (red) staining (**i**), and quantification of N-cadherin expression by integrated density (**ii**), n=26. **B)** qPCR data from hMSCs cultured on 2D PAAm hydrogels for 3 days showing fold-change in gene expression of N-cadherin (left) and β-catenin (right) relative to control with stronger elastic character (0.12 loss tangent samples) and normalised to GAPDH, n=3. **C)** Representative images of DAPI-stained hMSCs after 3 days culture in 3D PEG-MAL hydrogels. **D)** qPCR data from hMSCs cultured in 3D PEG-MAL hydrogels for 3 days showing fold-change in gene expression of N-cadherin (left) and β-catenin (right) relative to the control with stronger elastic character (0.1 loss tangent samples) and normalised to GAPDH, n=3. For all figures, data are represented as mean ±standard deviation and differences are considered significant for p≤ 0.05 using one-way ANOVAs and t-test for multiple and pairwise comparisons respectively (* p ≤ 0.05, ** p ≤ 0.01). All hydrogels were functionalised with 2 mM RGD peptide to allow cell adhesion. All PEG-MAL hydrogels were crosslinked using peptide ratios of 1% VPM and 99% scrambled VPM. Scale bars = 20 μm.

### hMSC chondrogenesis is facilitated in matrices with a higher loss tangent

Following the observations that the adhesive, mechanotransductive and cell-cell interactive behaviour of hMSCs in environments with a higher tan(*δ*) was representative of a chondrogenic phenotype, we next characterised their early and late chondrogenic differentiation in basal conditions. SOX9 is arguably the master regulator of chondrogenesis; its high expression is crucial for maintenance of the chondrocyte phenotype (33) and regulates the expression of cartilage matrix markers collagen II and aggrecan through direct binding and regulation of their promotor elements (34). Here, we observed an increase in SOX9 expression for hMSCs in environments with a higher tan(*δ*) both in 2D and in 3D (**Figures 5A, 5B and Supplementary Figure 9A**), and a concomitant downregulation in early osteogenic marker Runx2 expression (**Supplementary Figure 9B**). Runx2 is a master regulator of osteogenesis and one of the main antagonists of SOX9; there is a clear interdependent relationship between both proteins, as high Runx2 levels depress SOX9 expression (35), while elevated SOX9 inhibits Runx2 (36). Interestingly, we also observed an increased expression of Piezo1 for hMSCs in environments with a stronger viscous character (**Supplementary Figure 10)**; Piezo1, a mechanosensitive, stress-activated Ca^2+^ ion channel (37) recently shown to interact directly with cadherins (38), has been implicated as a marker of chondrogenesis, as it is robustly expressed in articular chondrocytes (39).

**Figure 5.**
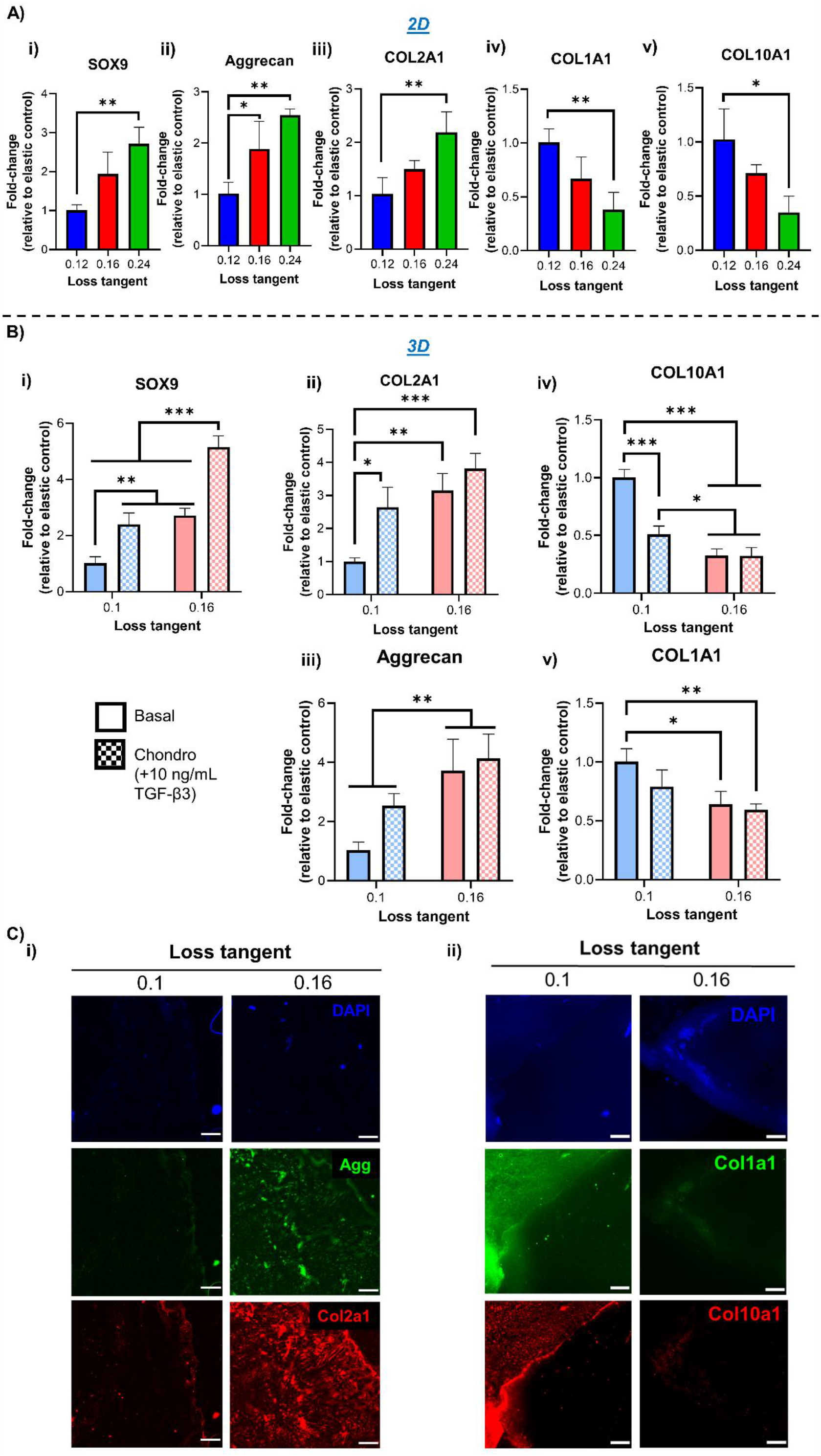
hMSC chondogenesis is promoted in matrices with a higher loss tangent. **A)** qPCR data showing fold-change in gene expression from hMSCs following 3-day culture on 2D PAAm hydrogels for SOX9 (**i**) and 7 days for Aggrecan (**ii**), COL2A1 (**iii**), COL1A1 (**iv**) and COL10A1 (**v**), n=3. **B)** qPCR data showing fold-change in gene expression from hMSCs cultured in 3D PEG-MAL hydrogels for 3 days for SOX9 (**i**) and 7 days for Aggrecan (**ii**), COL2A1 (**iii**), COL1A1 (**iv**) and COL10A1 (**v**), n=3. Cells were cultured in either basal medium or chondrogenic medium containing 10 ng/mL TGF-β3. **C)** Representative immunofluorescence images of hMSCs and surrounding matrix deposition following culture in 3D PEG-MAL hydrogels for 3 weeks and staining for DAPI (blue) with Aggrecan (green) and Col2a1 (red) (**i**) or with Col1a1 (green) and Col10a1 (red) (**ii**). For all figures, data are represented as mean ±standard deviation and differences are considered significant for p≤0.05 using oneway ANOVA for multiple comparisons (* p ≤ 0.05, ** p ≤ 0.01, *** p ≤ 0.001). All hydrogels were functionalised with 2 mM RGD peptide to allow cell adhesion. All qPCR data was made relative to control sample with stronger elastic character (0.12/0.1 loss tangent) and normalised to GAPDH. All PEG-MAL hydrogels were crosslinked using peptide ratios of 1% VPM and 99% scrambled VPM. Scale bars = 100 μm.

To further characterise the differentiation of hMSCs, we investigated the expression of cartilage matrix markers COL2A1 and aggrecan after prolonged culture times. We observed elevated expression of both markers in environments with higher tan(*δ*); this was accompanied by a downregulation in markers of chondrocyte hypertrophy COL10A1 and fibrocartilage COL1A1 compared to gels with a lower tan(*δ*) (**Figures 5A, 5B and Supplementary Figure 9C, D**). These observations also suggest that mature osteogenesis is favoured in more elastic environments. Indeed, collagen I is the most prominent organic component of bone and supports osteogenesis of stem cells cultured using collagen I matrices (40) while collagen X is involved in bone formation through endochondral ossification (41). Furthermore, the use of chondrogenic induction medium during culture enhanced the expression of cartilage matrix deposition only in more elastic environments, while it had no effect in higher tan(*δ*) hydrogels, suggesting that their mechanical properties alone saturated the cell response (**Figure 5B and Supplementary Figure 9C, D**). Matrix secretion was further characterised via staining of aggrecan and COL2A1 in PEG-MAL gels, showing elevated levels of secreted cartilage matrix in hydrogels with a higher tan(*δ*) compared to those with a lower one (**Figure 5C and Supplementary Figure 11**).

Overall, the higher loss tangent of the isoelastic hydrogels developed in this study influenced the adhesive, mechanotransductive, and cell-cell behaviour of hMSCs to favour commitment to a chondrogenic lineage. Importantly, the downregulation of chondrocyte hypertrophy and fibrocartilage markers suggests neocartilage formation, as evidenced by the matrix markers staining in 3D environments (**Figure 5C and Supplementary Figure 11**). Finally, it is important to highlight that the observed hMSC behaviour occurs independently of gel degradability. Indeed, when the amount of degradable crosslinker VPM was increased from 1 to 100%, similar responses were observed, in terms of adhesion, early signalling and matrix markers expression (**Supplementary Figure 12**).

## Discussion

Most studies investigating the role of stem cell mechanosensitivity in chondrogenesis focus solely on the elasticity of the cells’ environment, with the Young’s modulus of the substrates ranging from ~10 to ~1000 kPa (2) However, the viscous nature of the materials is generally disregarded. This has led to contradictory results, which could partly be attributed to interference from unreported variability in viscous modulus (2). While some studies have suggested that the viscous character of the substrates may be important during stem cell chondrogenesis, its contribution in the materials employed in these works is accompanied by a change in the elastic modulus or by the confounding effect of other biochemical cues (6–8). Therefore, the role of the materials’ viscous component in these biological processes remains elusive. Our work provides a comprehensive study of hMSC response to a variation of the viscous properties of 2D and 3D culture environments, eliminating any potential influence of the elastic character of the substrate. Using isoelastic matrices in growth factor-free conditions, we have explored a variety of cell responses to changes in substrate viscosity, including adhesion and spreading behaviour, mechanotransduction, cell-cell signalling, and differentiation.

Our results obtained on isoelastic matrices with Young’s moduli of ~12 kPa indicate that matrices exhibiting a more viscous character (high tan(*δ*)) promote a chondrogenic hMSC phenotype, facilitated by a rounded cell shape with reduced adhesion and low cytoskeletal tension (**Figure 6**). Instead, more elastic matrices (lower tan(*δ*)) support increased cell spreading and cytoskeletal tension, with high expression of integrins and focal adhesions that apply high traction forces; this promotes the expression of genes and proteins implicated in osteogenesis, fibrocartilage and chondrocyte hypertrophy (**Figure 6**). The reduced cell spreading observed in more viscous environments coincides with a decrease in nuclear mechanotransduction of YAP and in lamin A/C levels; YAP, which is considered a mechanical rheostat, was previously less understood in the context of hydrogel-driven stem cell chondrogenesis (28). Here, we show that the mechanically-driven downregulation of YAP nuclear translocation is correlated with a shift from an osteogenic to a chondrogenic response of hMSCs. Viscous matrices also lead to a downregulation of ROCK and Rac1 signalling, with ROCK being known to negatively regulate chondrogenesis (29, 42–44). Moreover, the matrix viscous component affects cell-cell contacts, with a high expression of N-cadherin and a downregulation of β-catenin in more viscous environments; such repression of Wnt signalling has been previously shown to stimulate early chondrogenesis (19, 20, 45). Collectively, these cellular responses facilitate stem cell chondrogenesis through increased expression of SOX9, COL2A1 and aggrecan in viscous matrices, while an osteogenic phenotype is favoured in more elastic matrices. Importantly, here we also demonstrate that the observed effects are independent of the material platform used and of its dimensionality. Similar viscoelastic properties of matrices with different composition prompt similar responses, either when cells are seeded on 2D PAAm, or when they are encapsulated within 3D PEG-MAL matrices. In the latter case, we critically show that the intrinsic viscous character of the matrix is the dominant factor in determining cell response rather than degradability, as higher VPM content does not alter the increased chondroinductive potential of viscous environments compared to their more elastic counterparts (**Supplementary Figure 12**).

In conclusion, we have shown that the cells’ mechanotransductive response to the viscous nature of their environment is key to determine the stem cell chondrogenic fate. Controlling the matrices’ viscous nature alone provides a growth factor-free, purely mechanically regulated way of efficiently targeting chondrogenesis of hMSCs and promoting the formation of neocartilage. The viscous and elastic components of hydrogels can therefore be better utilised as valuable parameters during the engineering of cartilage (or indeed other tissues), to harness the mechanosensitive response of stem cells and direct their fate towards specific lineages.

**Figure 6.**
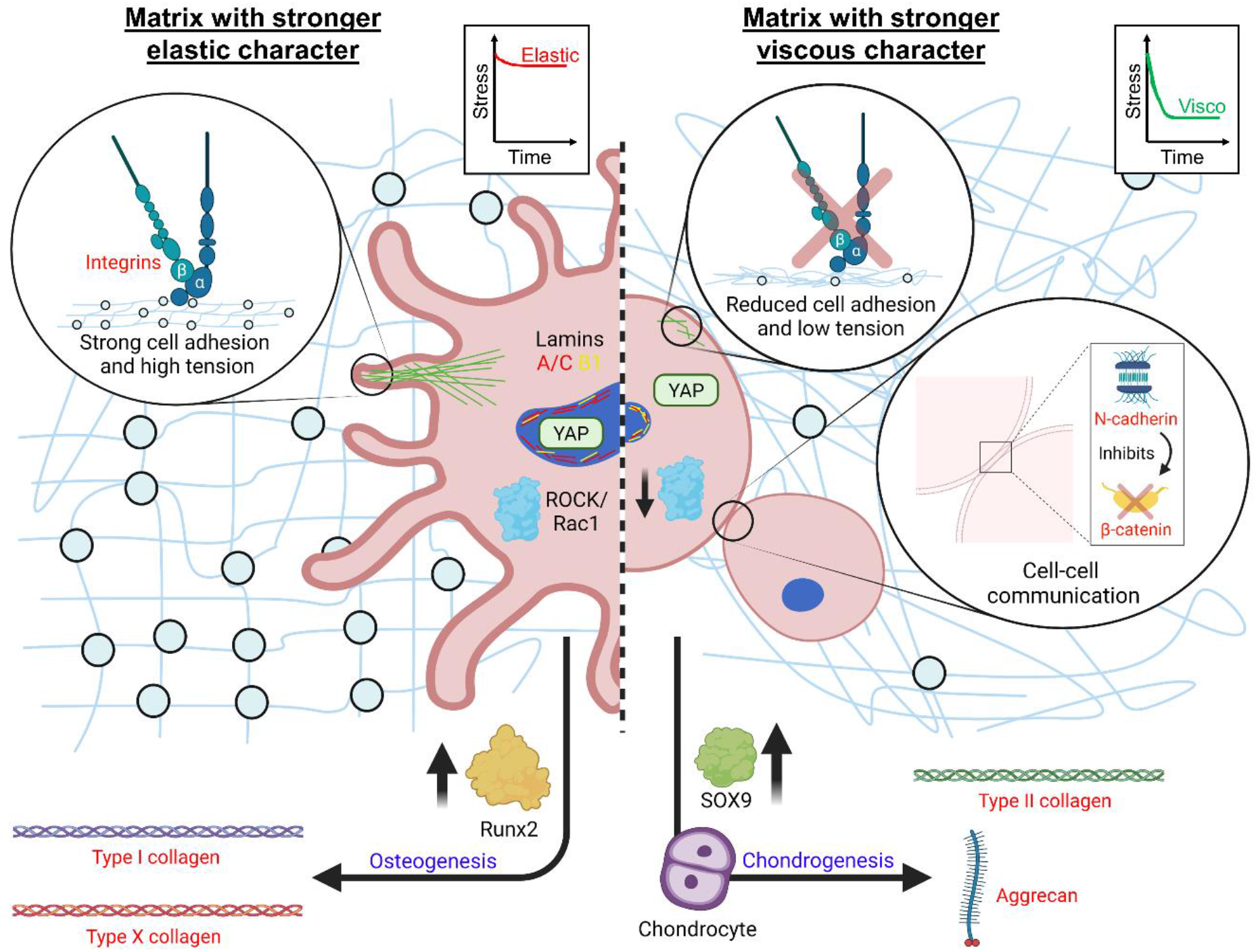
Mechanoregulation of hMSC differentiation in microenvironments with fixed elasticity and varied loss tangent. Using 12 kPa hydrogels, in an environment with a stronger elastic character hMSCs take on a highly spread, tensile phenotype with high expression of integrins and focal adhesions that apply high traction forces. This behaviour facilitates mechanotransduction and expression of genes and proteins implicated in osteogenesis, fibrocartilage and chondrocyte hypertrophy. In an environment with a higher viscous component, hMSCs take on a small, rounded phenotype with low cytoskeletal tension (comparable to a ROCK/Rac1-inhibited phenotype) and reduced integrin expression that diminishes cell traction forces to the ECM. This response inhibits mechanotransduction and promotes cell-cell contact through cell clustering, with increased N-cadherin expression and inhibition of β-catenin activity. Collectively, these responses drive chondrogenesis through increased expression of SOX9 and secretion of cartilage matrix markers. Figure generated using BioRender online software.

## Acknowledgements

This work was funded by a grant from the UK Regenerative Medicine Platform. MC and DG, respectively, acknowledge MRC funding (MR/S005412/1) and Royal Society of the United Kingdom funding under the Wolfson award (RSWF/FT/191020).

## Materials & Methods

### Peptides

**Table 1.** Names and sequences of peptides used (provided by GenScript).

| **Peptide name** | **Sequence** |
| --- | --- |
| RGD | GRGDSPC |
| FITC-RGD | GRGDSPC plus N-T:FITC-Ahx (N-Terminal) |
| VPM | GCRDVPMSMRGGDRCG |
| Scram VPM | GCRDVSPMGRMGDRCG |

### Antibodies (Abs)

**Table 2.** Names of primary Abs and suppliers used.

| **Abs** | **Supplier** | **Product code** | **Dilution** |
| --- | --- | --- | --- |
| SOX9 | Santa Cruz | sc-166505 | 1:250 |
| Lamin B1 | ProteinTech Group | 66095-1 | 1:250 |
| Lamin A/C | Santa Cruz | sc-376248 | 1:250 |
| Aggrecan | Santa Cruz | sc-67513 | 1:250 |
| COL2A1 | Santa Cruz | sc-518017 | 1:250 |
| COL1A1 | Santa Cruz | sc-59772 | 1:250 |
| COL10A1 | Abcam | ab58632 | 1:200 |
| YAP | Santa Cruz | sc-101199 | 1:250 |
| Runx2 | Santa Cruz | sc-390351 | 1:250 |
| N-cadherin | BD Biosciences | 610920 | 1:250 |
| p-FAK | Millipore | 05-1140 | 1:250 |
| Piezo1 | Novus Bio | NBP1-78537 | 1:50 |

**Table 3.** Names of secondary Abs and fluorophore-conjugated phalloidins used with their suppliers.

| **Abs** | **Supplier** | **Product code** | **Dilution** |
| --- | --- | --- | --- |
| Alexa Flour 488 | Thermo Fisher | A-11055/ A-11008 | 1:250 |
| Alexa Flour 488 phalloidin | Thermo Fisher | A12379 | 1:250 |
| Alexa Flour 647 phalloidin | Thermo Fisher | A30107 | 1:250 |
| Cy3-conjugated | Jackson ImmunoResearch | 111-165-003/715-165-150 | 1:300 |

### Primers

**Table 4.**
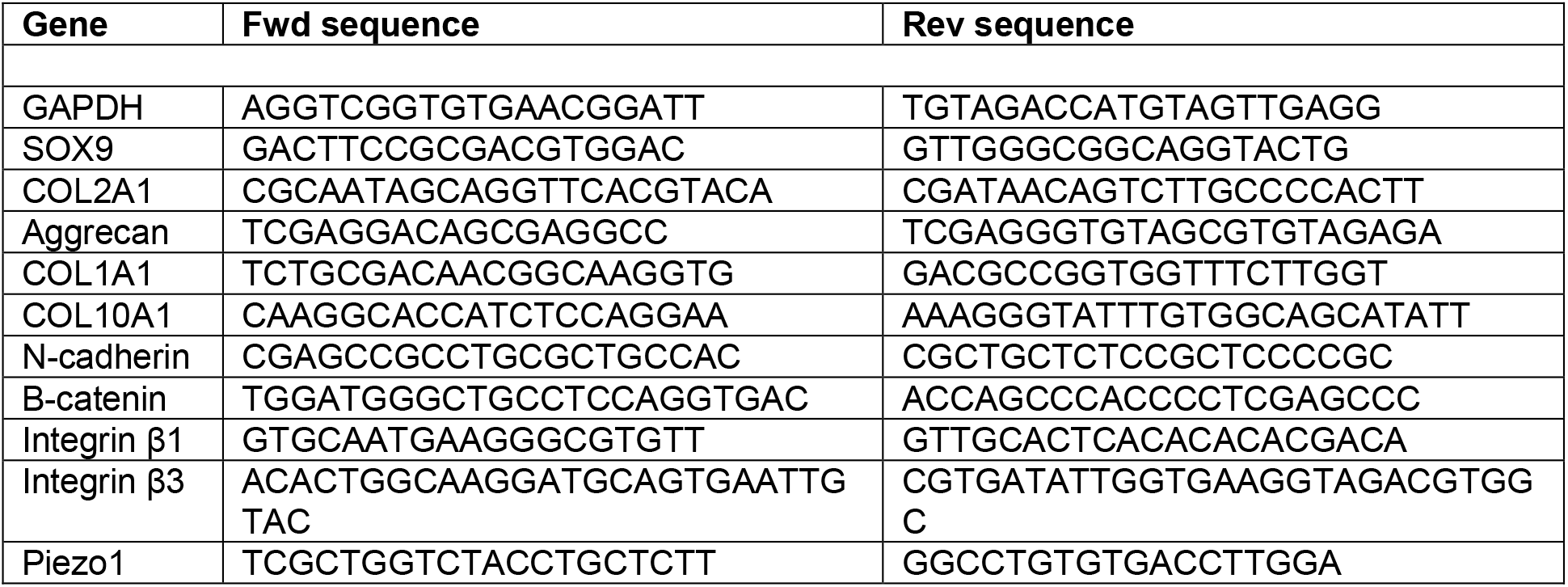
List of forward and reverse primers used for specific genes (provided by Thermo Fisher).

### Cell culture reagents

**Table 5.** List of cell culture reagents used for basal and chondrogenic culture of hMSCs.

| Reagent | Supplier/product |
| --- | --- |
| Dulbecco's Modified Eagle Medium (DMEM) | Sigma-Aldrich |
| Mesenchymal Stem Cell Growth Medium 2 | Promocell |
| Fetal Bovine Serum (FBS) | Gibco |
| GlutaMAX (100X) | Gibco |
| Sodium pyruvate (100 mM) | Sigma-Aldrich |
| MEM non-essential amino acids (100X) | Gibco |
| Penicillin/streptomycin (10,000 U/mL) | Sigma-Aldrich |
| Amphotericin B (250 ug/mL) | Gibco |
| Dexamethasone | Sigma-Aldrich |
| Insulin-transferrin-selenium (ITS) (100X) | Gibco |
| L-ascorbic acid | Sigma-Aldrich |
| L-proline | Sigma-Aldrich |
| TGF-β3 | R&D Biosystems/Biotechne |

### Glass preparation for 2D studies

12 mm glass coverslips and cover glass slides were RCA cleaned by washing in water and ethanol before heating for 10 mins at 65°C in a 5:1:1 solution of water:H2O2:NH3. After drying, cover glass slides were covered with Rain-X for 5 s, washed in ethanol and dried to use as hydrophobic glass slides. RCA-cleaned 12 mm coverslips were treated with specific silanes for either PAAm or PEG-MAL hydrogel fabrication in 2D. For PAAm hydrogels, coverslips were acryl-silanised submerged for 2 hrs in a 0.5% solution of 3-(Acryloyloxy)propyltrimethoxysilane (Alfa Aesar) in ethanol with 5% water. Coverslips were then dried and tempered by incubating for 1 hr at 120°C. For PEG-MAL hydrogels, coverslips were thiol-silanised by submerging for 3 hrs in a 10% solution of (3-Mercaptopropyl)trimethoxysilane (Sigma-Aldrich) in toluene before tempering by incubating at 100°C for 1 hr.

### 2D hydrogel fabrication

For PAAm hydrogels, all reagents were acquired from Sigma-Aldrich. Briefly, 1 mL volumes were prepared using stock solutions of 40% acrylamide and 2% N,N’-methylenebisacrylamide mixed in different ratios for specific gel compositions (**Table 6**). Solution volumes were then made up to 1 mL with milli-Q water, 2.5 μL tetramethylethylenediamine (TEMED) and 7.5 μL 10% ammonium persulfate (APS) and mixed thoroughly. 10 μL of solution was spotted onto hydrophobic glass slides before placing acrylsilanised glass coverslips onto the spots. Gelation was allowed to occur at room temperature for 30 min before detaching and swelling in water overnight at 4°C.

**Table 6.**
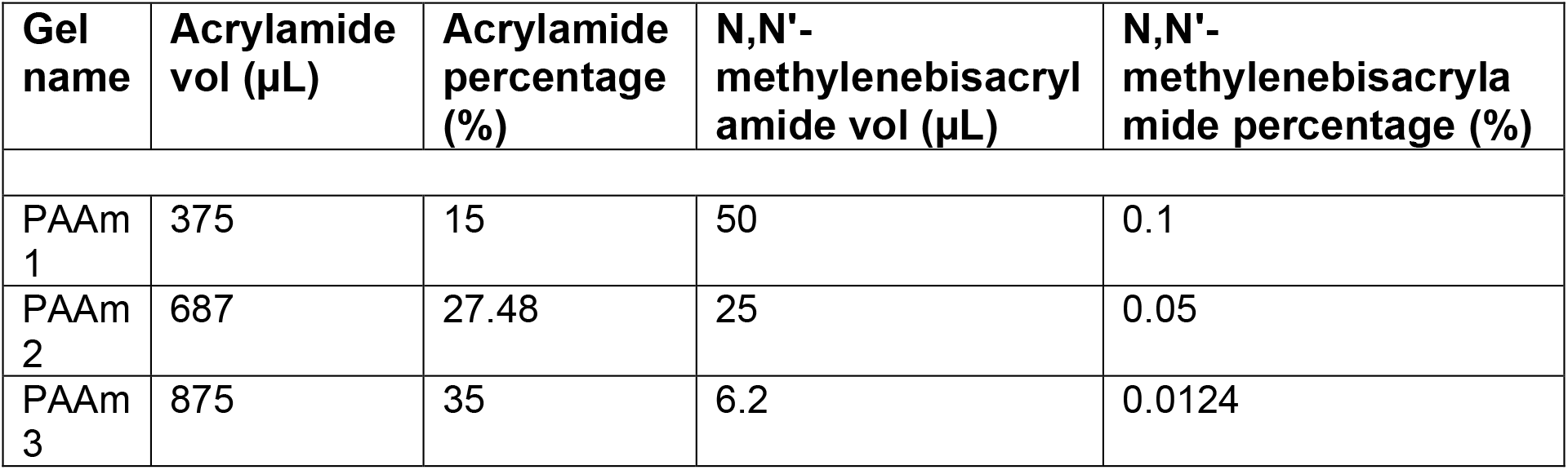
Acrylamide and N,N’-methylenebisacrylamide ratios for PAAm hydrogels.

| Gel name | Acrylamide vol (μL) | Acrylamide percentage (%) | N,N'-methylenebisacrylamide vol (μL) | N,N'-methylenebisacrylamide percentage (%) |
| --- | --- | --- | --- | --- |
| PAAm 1 | 375 | 15 | 50 | 0.1 |
| PAAm 2 | 687 | 27.48 | 25 | 0.05 |
| PAAm 3 | 875 | 35 | 6.2 | 0.0124 |

For PEG-MAL hydrogels, stock solutions of 20 kDa and 40 kDa 8-armed PEG-MAL (Creative PEGWorks), VPM and RGD peptides were prepared in phosphate buffered saline (PBS). Initially, appropriate amounts of PEG-MAL were mixed together with 2 mM RGD peptide for 1 hour at room temperature before adding VPM (calculated to stoichiometrically crosslink all remaining reactive groups of each hydrogel) and PBS to make up to 50 μL and mixing thoroughly (**Table 7**). 10 μL of solution was spotted onto hydrophobic glass slides before placing thiolsilanised glass coverslips onto the spots. Gelation was allowed to occur for 1 hr at 37°C before detaching and swelling in PBS overnight at 4°C.

**Table 7.** PEG-MAL and VPM peptide ratios for PEG hydrogels.

| Gel name | PEG-MAL conc. (mg/mL) | PEG-MAL percentage (%) | VPM conc. (mg/mL) | VPM percentage (%) |
| --- | --- | --- | --- | --- |
| PEG 1 | 213.5 (20 kDa) | 21.35 | 72.6 | 7.26 |
| PEG 2 | 427 (40 kDa) | 42.7 | 72.6 | 7.26 |

### Water absorption

Hydrogels were formed, weighed and immersed in milli-Q water/PBS, for PAAM/PEG-MAL hydrogels respectively, to swell overnight. After 24 hrs, the solvent was removed, the hydrated samples were weighed again and the amount of water absorbed was calculated using **Equation 1** as follows:

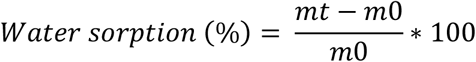

**Equation 1.** Swelling calculation for hydrogels; where *mt* is the weight of the hydrogel at a certain time and *m0* is the weight of the hydrogel after formation.

### Degradability

PEG-MAL hydrogels were formed, swollen overnight in PBS, and weighed prior to degradation. Then, a protease solution of 2.5 mg/mL collagenase D (Roche) in PBS was prepared and added to cover the samples before incubation at 37 °C. At each timepoint, all supernatant was removed by centrifugation at 4000 x g for 5 min and samples were weighed. Fresh protease solution was added at each timepoint and the degradation rate was calculated using **Equation 2** as follows:

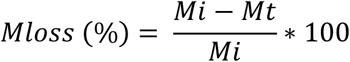

**Equation 2.** Degradability calculation for PEG-MAL hydrogels; where *Mloss* is the percentage of mass lost during degradation, *Mi* is the initial mass after swelling, and Mt is the mass at the different timepoints after the addition of the protease solution.

### Peptide functionalisation of PAAm hydrogels

PAAm gels prepared on coverslips were transferred to multiwell plates before covering with 0.2 mg/mL sulfosuccinimidyl 6-(4’-azido-2’-nitrophenylamino)hexanoate (sulfo-SANPAH) (Thermo Fisher) solution. Samples were placed in a 365 nm UV light source at a distance of ~3 inches and exposed for 10 min; this process was repeated 3 times. Gels were then washed with 50 mM HEPES buffer (pH 8.5) 3 times before covering with 2 mM RGD peptide solution (prepared in same HEPES buffer) and overnight incubation at 37°C. Gels were then washed with sterile-filtered milli-Q water to remove excess peptide.

### Atomic Force Microscopy

Using a NanoWizard 3 Bioscience AFM (JPK, Berlin, Germany), all measurements and cantilever calibrations were conducted at 37°C in aqueous environments (water/PBS for PAAM/PEG-MAL hydrogels respectively). Hydrogel samples were prepared on glass coverslips and superglued securely to tissue culture dishes before covering with liquid. Measurements and calibration of cantilever sensitivity against a stiff surface (tissue culture dish) and of spring constant, using the thermal noise method, were done using the JPK SPM software (version 6.1.192).

Force spectroscopy measurements were performed using a constant cantilever approach speed of 2.0 μm/s. ~0.3 N/m cantilevers (TL-CONT from Nanosensors) mounted with a 20 μm diameter spherical silica tip were used. Nanoindentation measurements were done using an indentation depth of ~1 μm. Microrheology measurements included a pause segment at constant height of 0.5 seconds after a ~1 μm indentation, followed by a 0.4 seconds oscillation/sine segment at a frequency of 10 Hz and amplitude of 10 nm; this was used to derive the viscoelastic response of the samples. From nanoindentation and microrheology measurements, the Young’s moduli and loss tangent were calculated using the Hertz model and microrheology processing functions of the JPK DP software (version 6.1.192). Force maps were carried out per sample condition to measure multiple points in different regions of the gels.

Quantitative AFM imaging was performed using ~0.3 N/m pyramidal, gold-coated cantilevers (PNP-TR-AU cantilevers from NanoWorld). Images were taken using 256 x 256 pixels within 20 um^2^ regions. Scans were performed at a frequency of 0.5 Hz using a setpoint of 4.5 nN and 2 μm z length.

### Rheology

Samples were prepared by forming hydrogels in PDMS moulds using 250 μL volumes and allowing to set before transferring to 6 well plates and swelling overnight at 4°C. For rheological measurements, samples were mounted onto a Physica MCR 301 rheometer (Anton Paar) and the linear viscoelastic region was determined by carrying out an amplitude sweep from 0.1 to 10% strain at 1 rad/s. Following this, a constant strain of 1% was used to obtain frequency sweeps from 0.5 to 50 rad/s.

### Stress relaxation measurements

Nanoindentation stress relaxation measurements were carried out using a nanoindentation device (Chiaro, Optics11 Life, Amsterdam, Netherlands) mounted on top of an inverted optical microscope (Zeiss Axiovert 200M, Zeiss). Measurements were performed following an adaptation of the protocol described in reference (46) using a cantilever with stiffness (k) of 0.52 Nm^-1^ holding a spherical tip of radius (R) of 27.5 μm.

Each gel was placed in a petri dish and stabilised with a drop of superglue between the silanised glass coverslip and the petri dish. All measurements were carried out at room temperature (~ 23°C) in milliQ water to maintain samples’ hydration. For each experimental condition, at least 100 indentations were performed, each spaced at least 100 μm from the previous. For each indentation, the probe moved at a strain rate of 5 μms^-1^ until it reached an indentation depth (δ) of 3 μm, which was maintained for 60 s using the instrument’s closed feedback Indentation control mode. This differs from most AFMs which maintain a constant height, resulting in an increasing indentation depth over the time of the experiment for viscoelastic materials (47).

Acquired data was cleaned using a previously published open-source software (available of GitHub, time branch of the project) (46). Briefly, the forward segment of the collected force-displacement (F-z) curves was inspected, and unsuccessful indentations discarded (i.e., indentations where contact was unsuccessful). Then, all segments were saved in a JSON file for further analysis. To analyse the stress relaxation behaviour of the material, a jupyter notebook was developed (https://github.com/GiuseppeCiccone96/stressrelaxnano). Briefly, force-time F(t) curves were first aligned to zero force if their baseline was negative. Then, the maximum of F(t) and its corresponding time was found, yielding the point (t0, F0). Curves were therefore aligned to 0 time by a horizontal shift equal to t0. Following this, the signal was cropped between t0 and the maximum time before retraction, i.e., only the part of the signal where the indentation was kept constant was retained. Following this, F(t) was normalised by dividing the whole signal by F0. Because individual curves were too noisy to be analysed, an average curve was found and used for quantification of the energy dissipation of the materials. Energy dissipation was quantified from the normalised signal using **Equation 3** as follows:

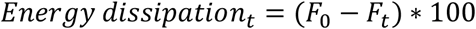

**Equation 3.** Energy dissipation calculation from normalised stress relaxation signal, where the subscript t denotes the maximum time in the averaged data.

### Cell Culture

Primary human mesenchymal stem cells (hMSCs) (Promocell) were thawed and resuspended in Mesenchymal Stem Cell Growth Medium 2 with supplement mix (Promocell) for expansion. Prior to seeding on or in the hydrogels, cells were serum-starved overnight in DMEM containing 1 % FBS. Cells were harvested by trypsinization at 70-80% confluency and cultured in a basal media of DMEM containing 10% FBS with biweekly media changes. High-glucose DMEM was used and supplemented with GlutaMAX (1X), sodium pyruvate (1 mM), non-essential amino acids (1X), penicillin/streptomycin (1%) and amphotericin B (2.5 μg/mL). Throughout all culturing, cells were incubated at 37°C in a 5% CO^2^ atmosphere.

For chondrogenic media cultures, modified DMEM was used containing 100 nM dexamethasone, 1X ITS, 50 μg/mL L-ascorbic acid-2-phosphate, 40 μg/mL L-proline and 10 ng/mL TGF-ß3.

### 2D cell seeding

For 2D cell studies, hydrogels on coverslips were sterilised under UV light for 30 min. Then, trypsinised cells were suspended in appropriate cell culture medium and seeded onto hydrogels at a density of 5,000 cells/cm^2^.

### 3D cell encapsulation in PEG-MAL hydrogels

For 3D cell studies, all PEG-MAL hydrogel reagents were sterilised under UV light for 30 min. Then, PEG-MAL was allowed to react with RGD peptide for 1 hr at room temperature while preparing cell pellets. Cell pellets were prepared by washing trypsinised cells in PBS via centrifugation (200 g for 5 min) to remove any presence of media components. Cell pellets were then suspended in the RGD-functionalised PEG-MAL solution at a density of 4 million cells/mL before adding VPM peptide and PBS to make up to 50 μl volumes. The full volume was pipetted into a multiwell plate and gelation was allowed to occur in a cell culture incubator for 1 hr. The cell-laden gels were then immersed in appropriate cell culture medium and allowed to recover in the incubator for 1 hr before replacing with fresh media for culturing.

### 3D sectioning

3D cultures were fixed using 4% formaldehyde for 30 min at 4°C before exchanging with 30% sucrose in PBS overnight at 4°C. Samples were then included in optimal cutting temperature compound (OCT) in cryomoulds and frozen using liquid nitrogen before storage at −80°C. Samples were cut using a cryotome in 20 μm sections and frozen at −80°C prior to immunostaining.

### Immunostaining

2D samples were washed with PBS before fixing with 4% formaldehyde for 30 min at 4°C. 3D samples were sectioned after fixing as explained above. Then, PBS washes were done followed by permeabilization in 0.1% Triton X-100 for 5 min at room temperature. Samples were then washed with PBS and blocked for 1 hr in 1% bovine serum albumin (BSA). After blocking, all primary Ab solutions were prepared in 1% BSA at appropriate dilutions (**Table 2**) and added to cover the samples before incubating overnight at 4°C. After primary Ab incubations, samples were washed 3 times with 0.5% Tween-20 prior to addition of secondary Ab solutions (diluted appropriately in 1% BSA, **Table 3**) and incubation at room temperature in the dark for 1 hr. Samples were then washed 3 times with 0.5% Tween-20 before mounting onto glass slides using VECTASHEILD antifade mounting medium with DAPI (Vector Laboratories). Visualisation was done using a fluorescence microscope (Zeiss AxioObserver Z1).

### Image analysis

Image processing was done using ImageJ (version 1.53t). Cells were measured by binarising nucleus and/or actin cytoskeleton in images using a threshold function. Then, the wand tracing tool was used to select the outline of the thresholded areas and the measure function was used to calculate morphological parameters, such as cell area and circularity; protein expression levels were measured via integrated density (normalised to cytoskeletal or nuclear area depending on region of expression within the cell). YAP expression was represented as a nuclear/cytoplasmic ratio; this was performed by measuring nuclear and cytoplasmic YAP expression independently and calculated as follows:

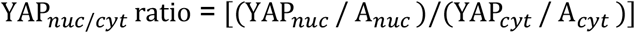

**Equation 4.** YAP’s integrated density fluorescence nucleus/cytoplasm ratio; where YAP_nuc_ is the integrated density of YAP in the nucleus, Anuc is the area of the nucleus, YAPcyt is the integrated density of YAP in the cytoplasm (**Equation 5**), and Acyt is the area of the cell cytoplasm (**Equation 6**).

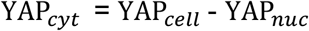

**Equation 5.** YAP’s integrated density fluorescence in the cytoplasm; where YAPcell is the integrated density of YAP in the entire cell

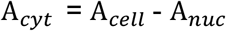

**Equation 6.** Definition of cytoplasmic area; where Ace⊓ is the area of the entire cell.

Focal adhesion analysis was performed on p-FAK stained samples using a previously described step-by-step method (48) and implementing it in ImageJ. Actin fibre anisotropy was performed using the FibrilTool plug-in as previously described (49).

### Pharmacological inhibition

Pharmacological inhibition of Rac1 and ROCK activity was achieved using 50 μM NSC-23766 (Tocris Bioscience) and 10 μM Y-27632 (Calibiochem) respectively. Inhibitors were added to cell cultures 1 hour prior to fixing.

### Traction force microscopy

Carboxylate-modified 0.2 μm FluoSpheres (Life Technologies) were prepared by sonicating the stock for 10 mins. This was then diluted 1:30 in milli-Q water and further sonicated for 15 mins. Immediately after sonication, 1:25 of bead dilution was incorporated into the gel recipes for PAAm hydrogels on coverslips before functionalisation with 2 mM RGD peptide (as previously described). Cells were seeded on the 2D gel surfaces (as described previously) and allowed to adhere for 24 hrs. Using an EVOS FL Auto microscope (Life Technologies), Z-stack images were taken through the cells and gels before and after trypsinisation. Cell traction forces were determined using ImageJ software by tracking the displacement of the FluoSpheres and then reconstructing the force field from the displacement data using the iterative particle image velocimetry (PIV) and FTTC plugins respectively (50)

### qPCR

All reagents were provided by QIAGEN unless otherwise stated. Cell lysis and RNA extraction was performed using a RNeasy mini kit. For 2D samples, cells were directly lysed on the surface of the gels before RNA extraction. For 3D samples, cells were released by incubating gels in a cell culture incubator in a protease solution of 2.5 mg/mL collagenase D (Roche) in PBS until gels were fully degraded. Cells were pelleted via centrifugation (200 g for 5 min) and then lysed for RNA extraction using an RNeasy Micro Kit. RNA quantity and purity were measured using a NanoDrop 1000 (Thermo Scientific) before performing cDNA synthesis using a QuantiTect Reverse Transcription kit. Real-time qPCR was performed using a model 7500 real-time PCR machine (Applied Biosciences) using SYBR green reagents from a QuantiFast SYBR Green PCR kit. 4 ng cDNA was used per gene, primers were used at 1 μM concentrations and GAPDH was used throughout as a housekeeping gene for normalisation of fold-changes in gene expression.

### Cell viability

Cells were encapsulated in PEG-MAL hydrogels and cultured in basal media. After 24 hrs, CCK-8 solution (Sigma-Aldrich) was pre-warmed for 5 mins at 37°C, added into the culture medium at a 10% concentration, and incubated for 4 hrs in a cell culture incubator. The solution was then removed, transferred to a multiwell plate, and measured at an absorbance of 450 nm by a NanoQuant Infinite M200 Pro plate reader (Tecan). Fresh media was added to the samples and the process was repeated after a further 24 hrs of culture.

### Statistical analysis

Data were analysed using GraphPad Prism software where normality tests were performed to determine whether to select parametric or non-parametric tests. Then appropriate one-wayANOVA or t-tests, for multiple or pairwise comparisons respectively, were used and differences were considered significant for p≤0.05 (* p ≤ 0.05, ** p ≤ 0.01, *** p ≤ 0.001).

