## Supplementary Information for "Mind the viscous modulus: The mechanotransductive response to the viscous nature of isoelastic matrices regulates stem cell chondrogenesis"

### Supplementary figures

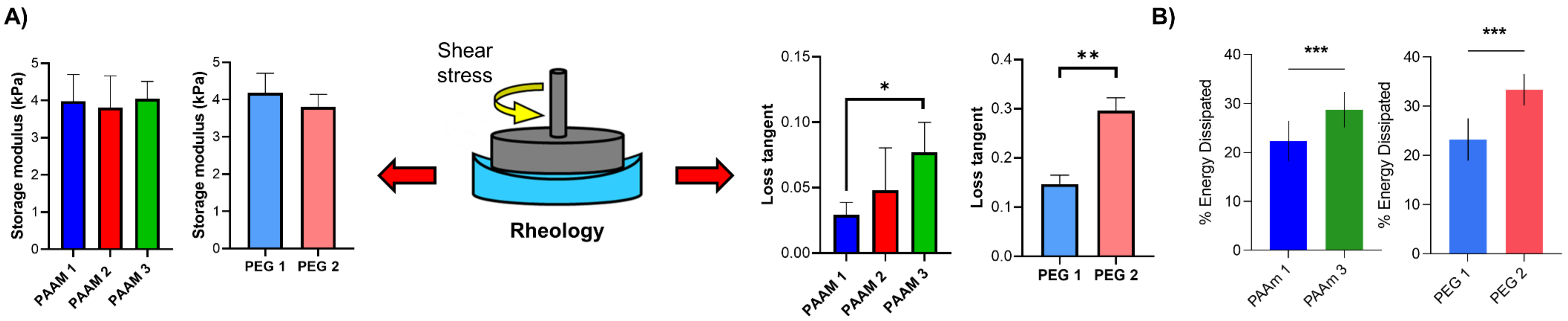


**Supplementary Figure 1. Rheological bulk characterisation shows comparable mechanical values to micro-scale AFM results and energy dissipation during stress relaxation is greater for gels with more viscous character. A)** Rheology-scale measurements of PAAm and PEG-MAL gels (graphical representation in middle) for storage moduli (left) and loss tangent (right) using linear region of frequency sweeps (50-0.5 rad/s) at 1% strain, n=3. **B)** Energy dissipation quantified from the normalised signal of stress relaxation measurements over 60 s for PAAm (left) and PEG (right) hydrogels, n=100. For all figures, data are represented as mean ± standard deviation and differences are considered significant for $p \leq 0.05$ using one-way ANOVAs or t-tests for multiple or pairwise comparisons respectively (* $p \leq 0.05$, ** $p \leq 0.01$, *** $p \leq 0.001$).

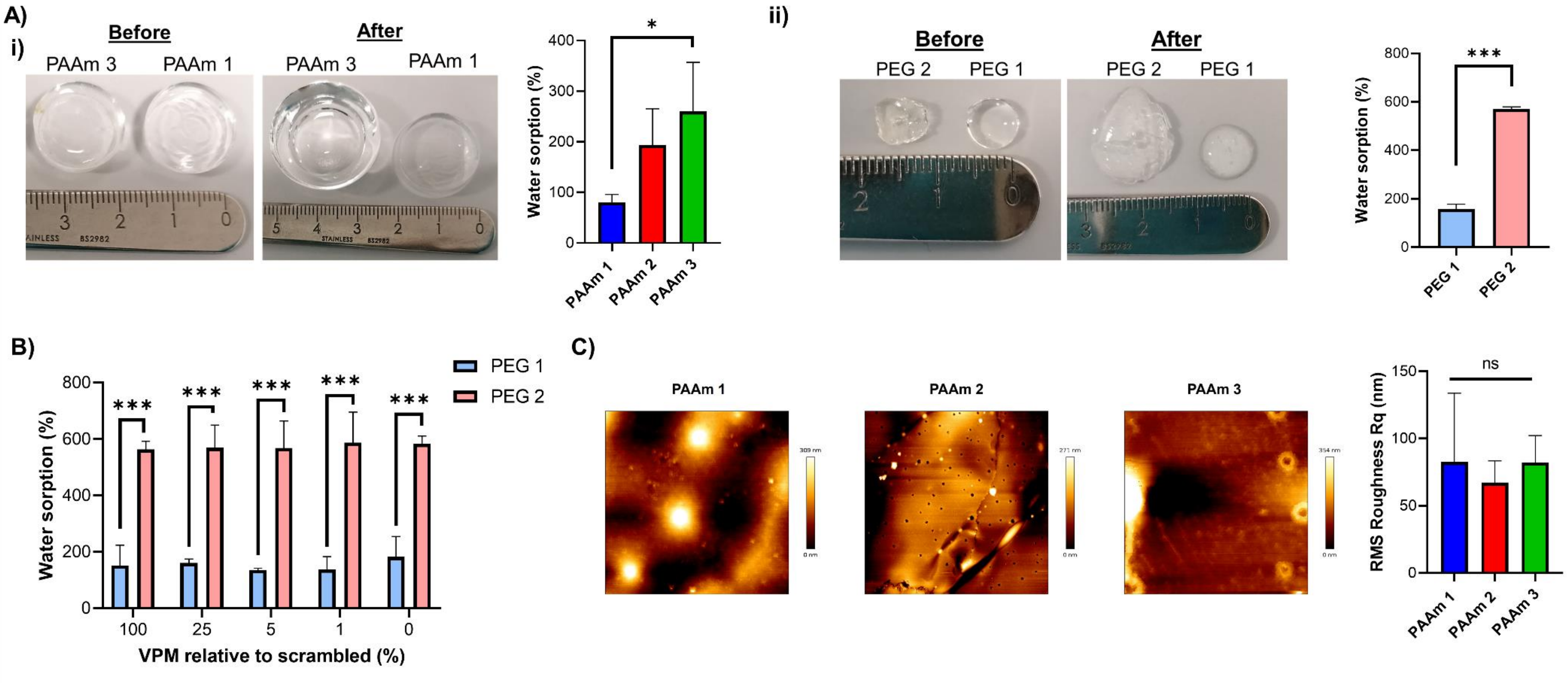


**Supplementary Figure 2. Swelling behaviour of PAAm and PEG-MAL hydrogels and topographical properties of PAAm hydrogels. A)** Swelling behaviour of PAAm gels following overnight incubation in water at 4ºC (**i**) with representative before/after images (left) and water sorption quantification (right), n=3. Swelling behaviour of PEG-MAL gels following overnight incubation in PBS at 4ºC (**ii**) with representative before/after images (left) and water sorption quantification (right), n=3. **B)** Water sorption quantification of PEG-MAL gels with different degrees of degradability, n=3. **C)** Representative images of PAAm hydrogel topography from height measurements during quantitative imaging mode of AFM (left) and quantification of RMS roughness rq, n=3). For all figures, data are represented as mean ± standard deviation and differences are considered significant for $p \leq 0.05$ using one-way ANOVAs or t-tests for multiple or pairwise comparisons respectively (* $p \leq 0.05$, *** $p \leq 0.001$).

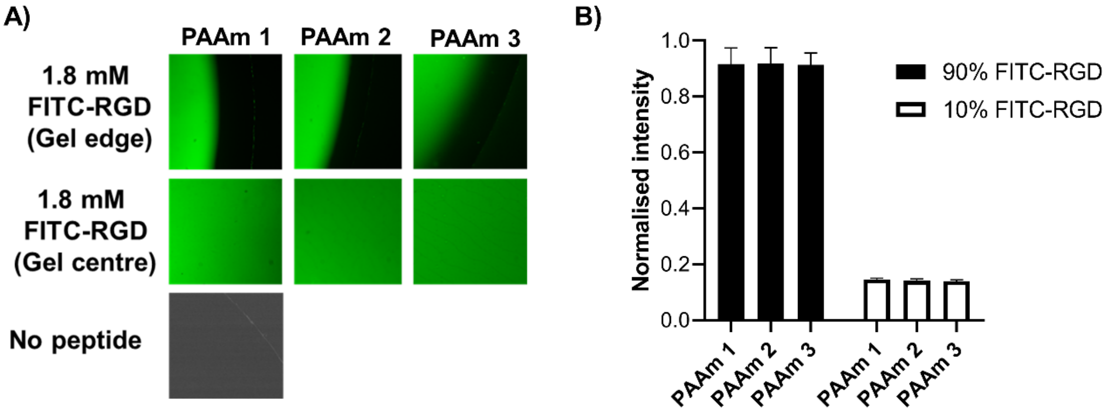


**Supplementary Figure 3. RGD Peptide functionalisation is comparable between PAAm hydrogels of different viscoelasticity. A)** Representative fluorescent microscopy images for 1.8 mM FITC-RGD functionalisation of viscoelastic PAAm hydrogels at edges and centre of gels. **B)** Normalised quantification of FITC-RGD intensities on PAAm hydrogel surfaces using 90% and 10% ligand densities of 2 mM FITC-RGD, n=10-13.

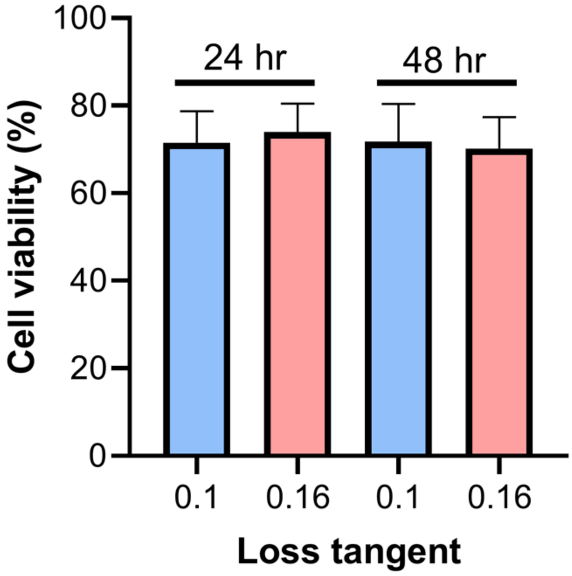


**Supplementary Figure 4. hMSC viability in PEG-MAL gels:** CCK-8 assays of hMSCs cultured in PEG-MAL hydrogels over 48 hours (relative to viability of known cell numbers in well plates). All hydrogels were functionalised with 2 mM RGD peptide to allow cell adhesion and were crosslinked using peptide ratios of 1% VPM and 99% scrambled VPM, n=9

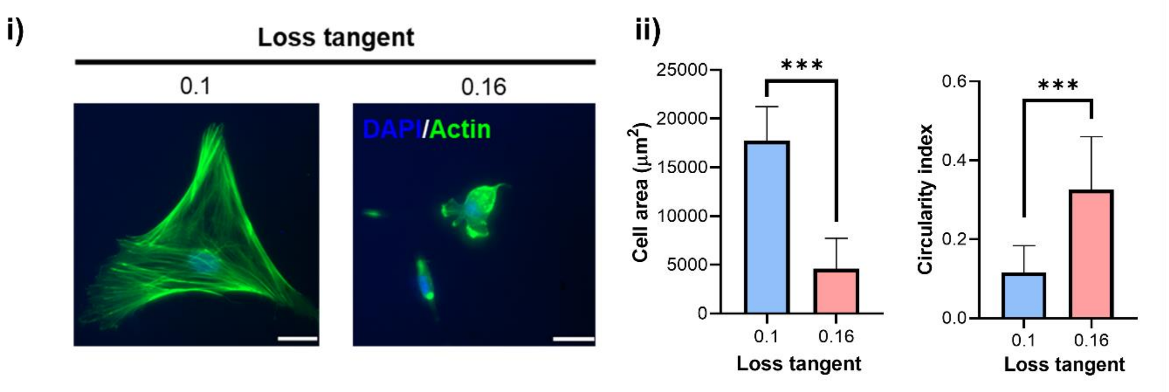


**Supplementary Figure 5. hMSC spreading behaviour on PEG-MAL hydrogels in 2D is comparable with that seen on PAAm gels.** Representative immunofluorescence images of hMSCs cultured for 3 days on PEG-MAL hydrogels with DAPI (blue) and actin staining (green) (**i**) with quantification of cell area (left) and circularity (right), n=15-24 (**ii**). For all figures, data are represented as mean ± standard deviation and differences are considered significant for $p \leq 0.05$ using t-tests for pairwise comparisons (*** $p \leq 0.001$). All hydrogels functionalised with 2 mM RGD peptide to allow cell adhesion. PEG-MAL hydrogels were crosslinked using 100% VPM peptide. Scale bar = 20 µm.

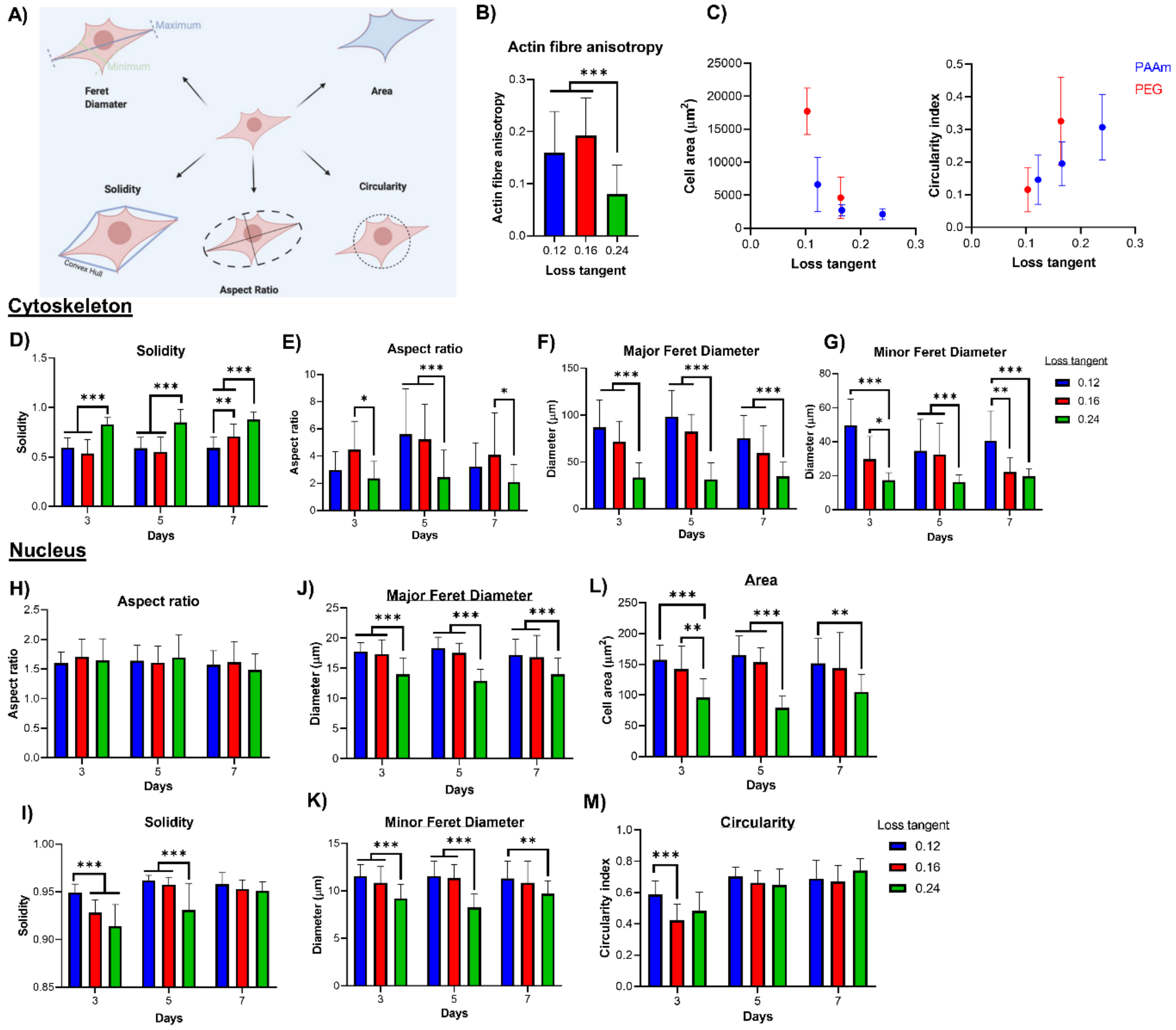


**Supplementary Figure 6. Cytoskeletal and nuclear morphological descriptors of hMSCs in viscoelastic environments over time. A)** Representation of shape descriptors: Feret diameters describe the longest (major) and shortest (minor) lengths between two parallel lines running perpendicular to those directions and tangent to opposite sides of the cell. The area describes the total two-dimensional size of the cell/nucleus. The solidity is the ratio between the area and the convex hull area and describes how convex the cell/nucleus is (1 for a convex cell without concavities). The circularity describes how close to a circle of the same area a cell/nucleus is (1 for a perfect circle). The aspect ratio describes the ratio of major/minor axis of an ellipse fitting the cell. **B)** Quantitative analysis of actin cytoskeleton organisation of hMSCs on PAAm hydrogels after 5 days; anisotropy of actin fibres calculated from immunofluorescence images using FibrilTool n=22-27. **C)** Comparative cell area (left) and circularity (right) for hMSCs cultured for 24 hours on PAAm and PEG-MAL hydrogels as a function of loss tangent, n=17-35. **D-G)** Cytoskeletal morphological descriptors of solidity (**D**), aspect ratio (**E**), major Feret diameter (**F**) and minor Feret diameter (**G**) for hMSCs on PAAm hydrogels over 7 days, n=10-32. **H-M)** Nuclear morphological descriptors of aspect ratio (**H**), solidity (**I**), major Feret diameter (**J**), minor Feret diameter (**K**), area (**L**) and circularity (**M**) for hMSCs on PAAm hydrogels over 7 days, n=19-123. For all figures, data are represented as mean ± standard deviation and differences are considered significant for $p \leq 0.05$ using one-way ANOVAs (* $p \leq 0.05$, ** $p \leq 0.01$, *** $p \leq 0.001$). All hydrogels functionalised with 2 mM RGD peptide to allow cell adhesion. PEG-MAL hydrogels were crosslinked using 100% VPM peptide.

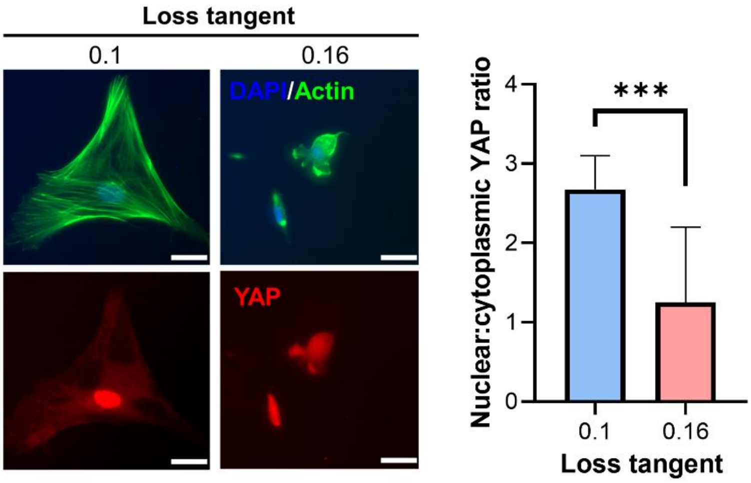


**Supplementary Figure 7. YAP mechanotransduction on PEG-MAL hydrogels in 2D is comparable with that seen on PAAm gels.** Representative immunofluorescence images of hMSCs cultured for 3 days on PEG-MAL hydrogels with DAPI, actin and YAP staining (left) and quantification of nuclear:cytoplasmic YAP ratio (right) for PAAm, n=17-23. Data are represented as mean ± standard deviation and differences are considered significant for $p \leq 0.05$ using t-tests (*** $p \leq 0.001$). All hydrogels functionalised with 2 mM RGD peptide to allow cell adhesion. PEG-MAL hydrogels were crosslinked using 100% VPM peptide. Scale bar = 20 µm.

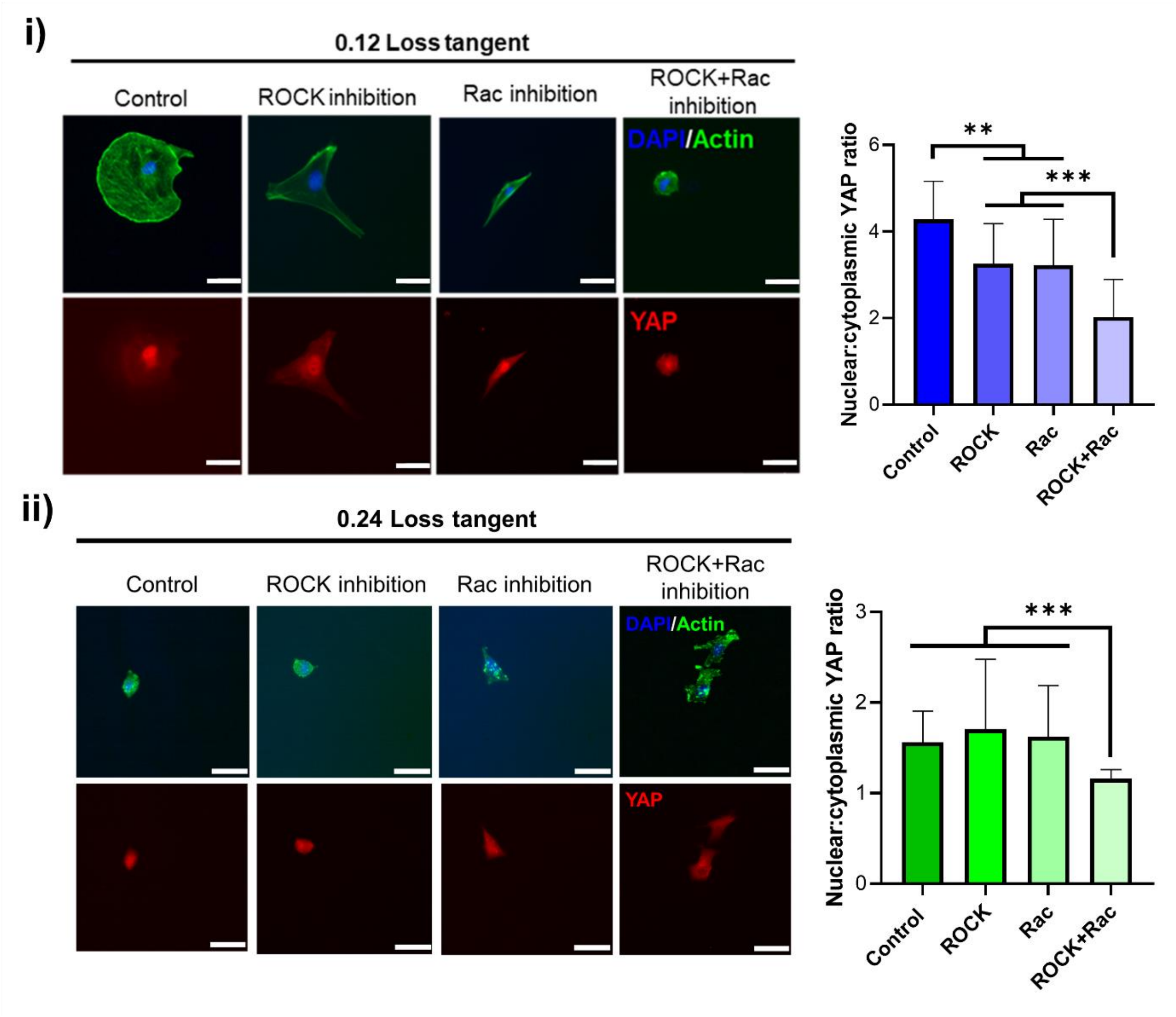


**Supplementary Figure 8. Disruption of hMSC cytoskeletal organisation on 2D PAAm hydrogels by ROCK and Rac1 inhibition reduces nuclear YAP on gels with stronger elastic character when inhibited individually or collectively, but requires collective inhibition on more viscous gels.** Representative immunofluorescence images of hMSCs cultured for 3 days, before 1 hour incubation with ROCK and Rac inhibitors, on 0.12 loss tangent (**i**) or 0.24 loss tangent (**ii**) PAAm hydrogels with DAPI (blue), actin (green) and YAP staining (red) (left) and quantification of nuclear:cytoplasmic YAP ratio (right) n=29-32. For all figures, data are represented as mean ± standard deviation and differences are considered significant for $p \leq 0.05$ using one-way ANOVAs (** $p \leq 0.01$, *** $p \leq 0.001$). All hydrogels functionalised with 2 mM RGD peptide to allow cell adhesion. Scale bars = 20 µm.

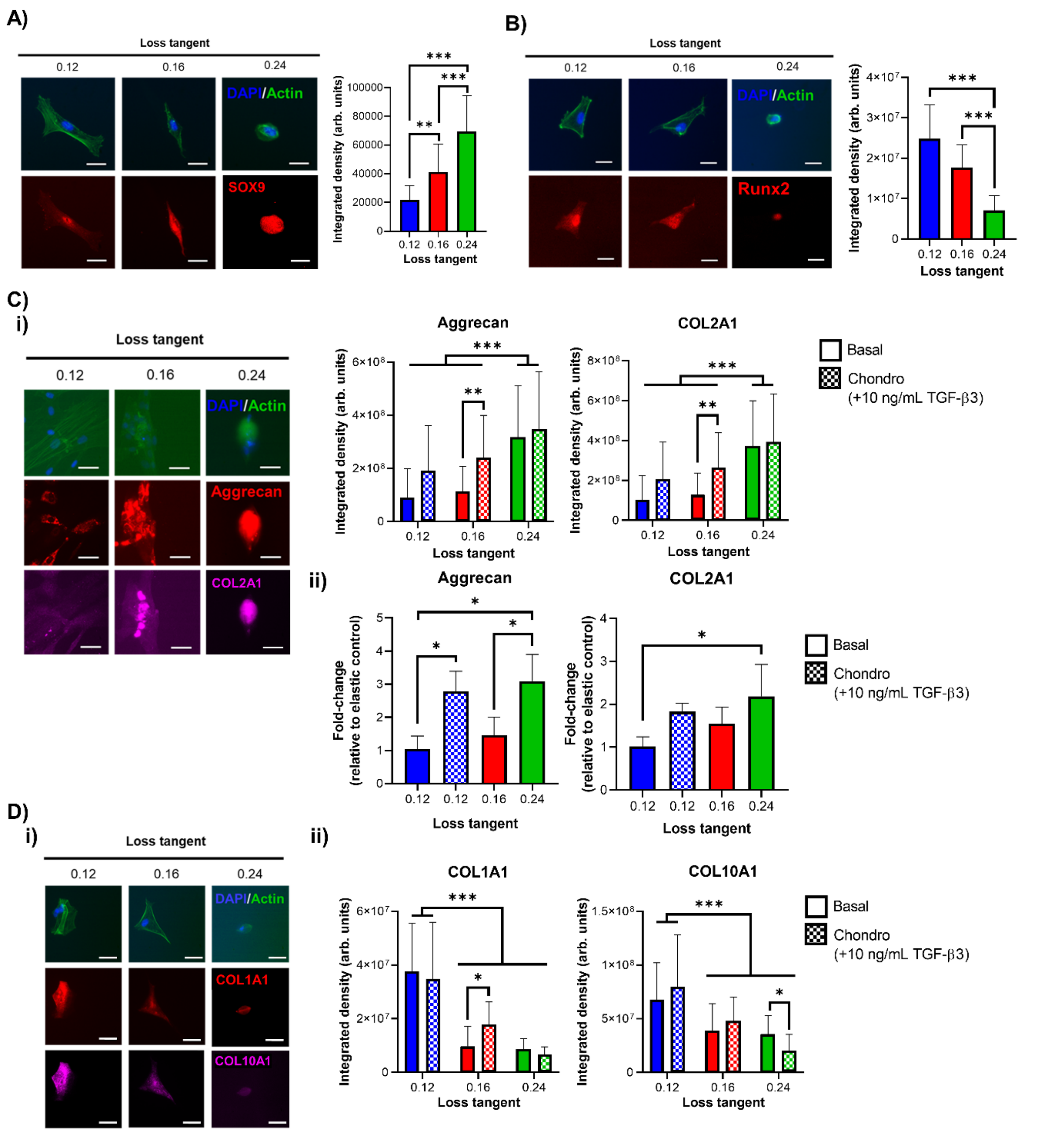


**Supplementary Figure 9. hMSC chondrogenesis and inhibition of fibrocartilage/chondrocyte hypertrophy is modulated by loss tangent in 2D. A)** Representative immunofluorescence images of hMSCs cultured for 7 days on viscoelastic PAAm hydrogels with DAPI (blue), actin (green) and SOX9 (red) staining (left) and quantification of SOX9 expression by integrated density (right), n=25-33 **B)** Representative immunofluorescence images of hMSCs cultured for 7 days on viscoelastic PAAm hydrogels with DAPI (blue), actin (green) and Runx2 (red) staining (left) and quantification of Runx2 expression by integrated density (right), n=26-29 **C)** Representative immunofluorescence images of hMSCs cultured for 3 weeks on viscoelastic PAAm hydrogels with DAPI (blue), actin (green), aggrecan (red) and COL2A1 (magenta) staining (**i**) and quantification of aggrecan (left) and COL2A1 (right) expression by integrated density in basal or chondrogenic media, n=22-32. (**ii**) qPCR data from hMSCs cultured on PAAm hydrogels for 7 days in basal or chondrogenic media showing fold-change in gene expression of aggrecan (left) and COL2A1 (right) relative to control with lower loss tangent (0.12 loss tangent basal samples) and normalised to GAPDH, n=3. **D)** Representative immunofluorescence images of hMSCs cultured for 3 weeks on viscoelastic PAAm hydrogels with DAPI (blue), actin (green), COL1A1 (red) and COL10A1 (magenta) staining (**i**) and quantification of COL1A1 (left) and COL10A1 (right) expression by integrated density in basal or chondrogenic media, n=21-32 (**ii**). For all figures: data are represented as mean ± standard deviation and differences are considered significant for $p \leq 0.05$ using one-way ANOVA for multiple comparisons (* $p \leq 0.05$, ** $p \leq 0.01$, *** $p \leq 0.001$); all hydrogels functionalised with 2 mM RGD peptide to allow cell adhesion. Scale bars = 20 µm.

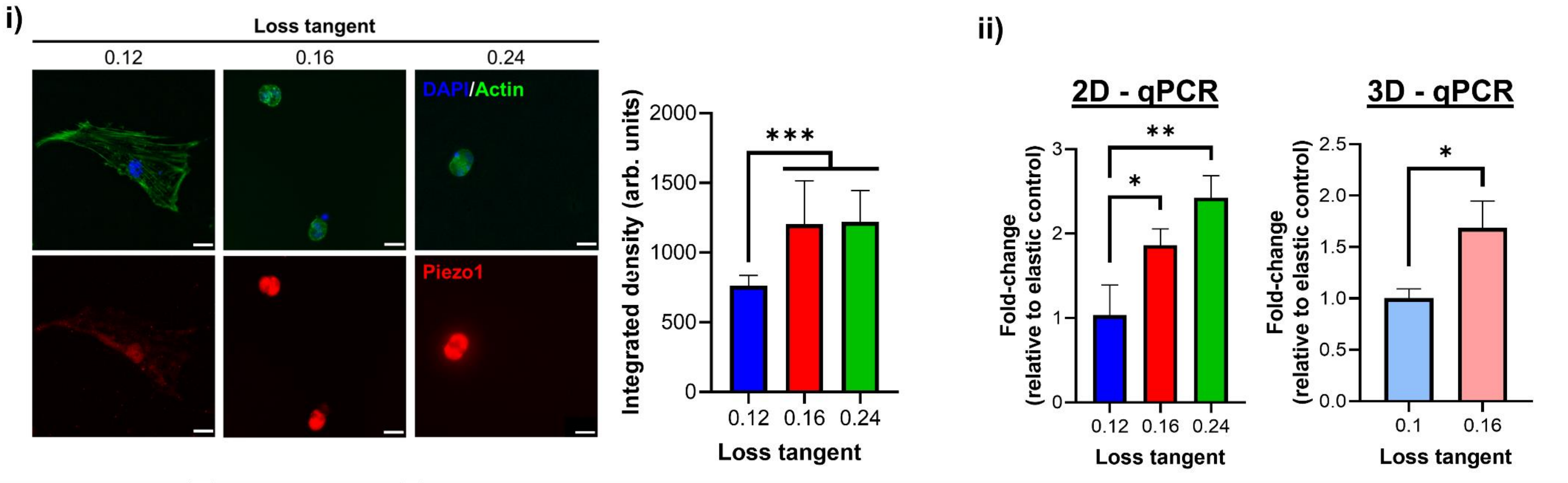


**Supplementary Figure 10. Piezo1 expression increases for hMSCs cultured using hydrogels with higher loss tangent.** hMSCs cultured for 3 days on PAAm gels before staining for DAPI (blue), actin (green), and Piezo1 (red) **(i)** with representative images (left) and Piezo1 integrated density quantification, (right) n=25-33. qPCR data from hMSCs cultured using hydrogels for 3 days showing fold-change in gene expression of Piezo1 **(ii)** using PAAm (left) or PEG (right) hydrogels, n=3. For all figures, data are represented as mean ± standard deviation and differences are considered significant for $p \leq 0.05$ using one-way ANOVAs or t-tests for multiple or pairwise comparisons respectively (* $p \leq 0.05$, ** $p \leq 0.01$, *** $p \leq 0.001$). All hydrogels functionalised with 2 mM RGD peptide to allow cell adhesion. All qPCR data was made relative to control sample with stronger elastic character (0.12/0.1 loss tangent) and normalised to GAPDH. PEG-MAL hydrogels were crosslinked using 100% VPM peptide. Scale bar = 20 µm.

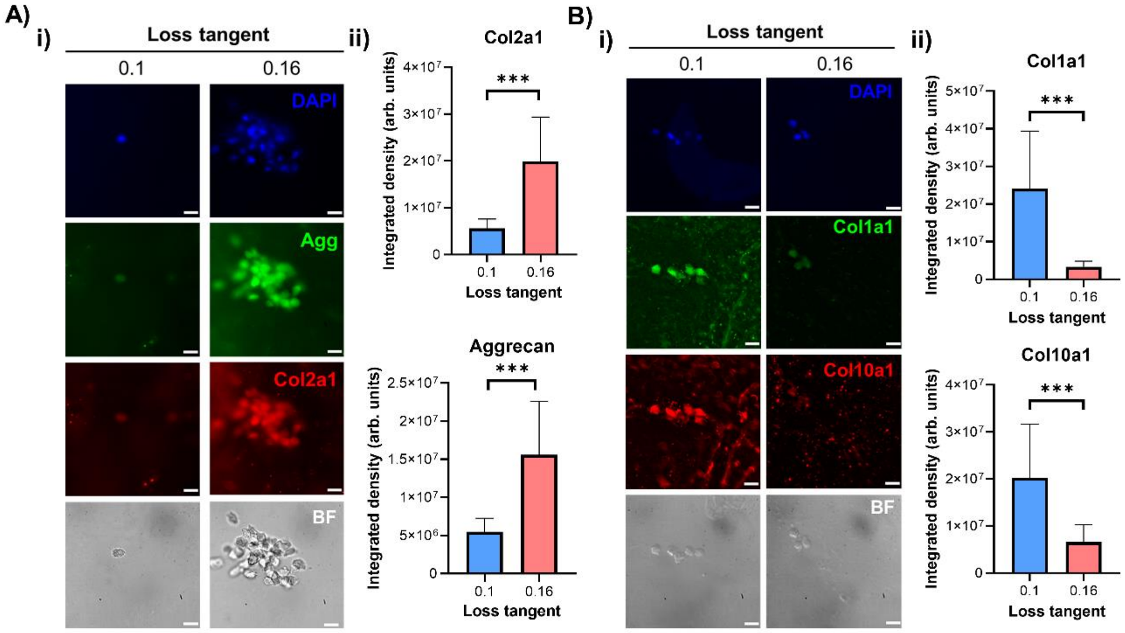


**Supplementary Figure 11. Cell expression of cartilage matrix markers is upregulated and fibrocartilage/hypertrophy markers are downregulated in 3D hydrogels with higher loss tangent. A)** Representative immunofluorescence and bright field images of hMSCs cultured for 3 weeks in PEG-MAL hydrogels stained with DAPI, aggrecan and Col2a1 (**i**) with quantification for expression using integrated density (**ii**), n=44-52. **B)** Representative immunofluorescence and bright field images of hMSCs cultured for 3 weeks in PEG-MAL hydrogels stained with DAPI, Col1a1 and Col10a1 (**i**) with quantification for expression using integrated density (**ii**), n=46-47. For all figures, data are represented as mean ± standard deviation and differences are considered significant for $p \leq 0.05$ using t-tests for pairwise comparisons (*** $p \leq 0.001$). All hydrogels were functionalised with 2 mM RGD peptide to allow cell adhesion. All PEG-MAL hydrogels were crosslinked using peptide ratios of 1% VPM and 99% scrambled VPM. Scale bars = 20 µm.

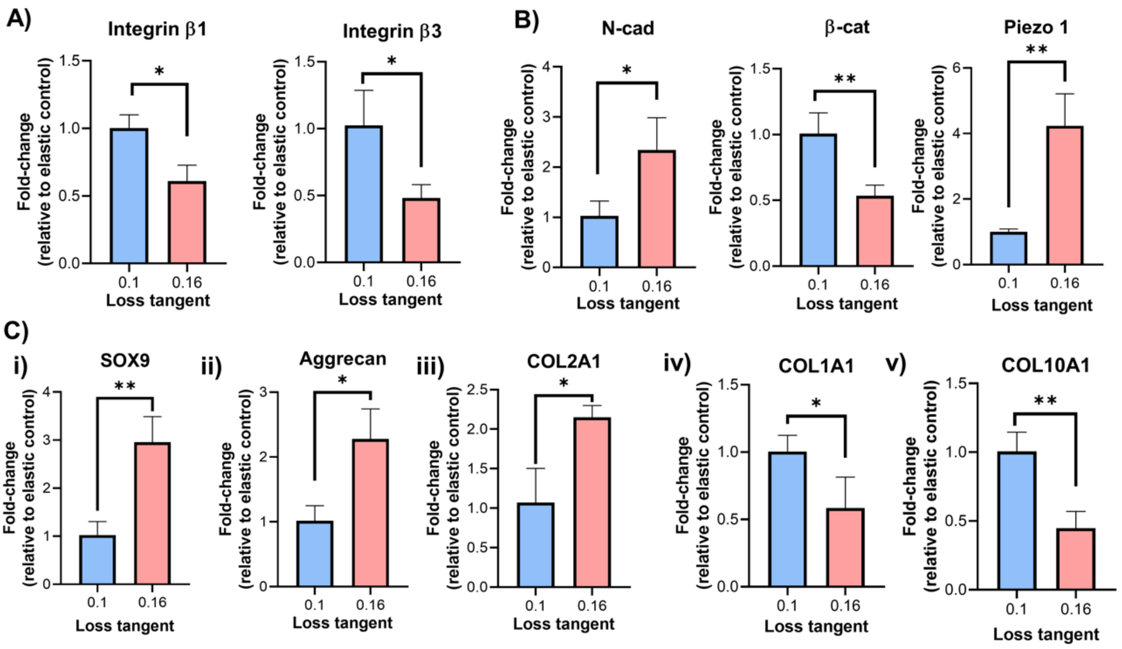


**Supplementary Figure 12. Independently of hydrogel degradability, matrix loss tangent regulates the expression of genes involved in adhesion, mechanotransduction, and chondrogenic differentiation. A)** qPCR data from hMSCs cultured in PEG-MAL hydrogels for 3 days showing fold-change in gene expression of integrin β1 (left) and integrin β3 (right), n=3. **B)** qPCR data from hMSCs cultured in PEG-MAL hydrogels for 3 days showing fold-change in gene expression of N-cadherin (left), β-catenin (middle) and Piezo1 (right), n=3. **C)** qPCR data showing fold-change in gene expression from hMSCs cultured in PEG-MAL hydrogels for 3 days for SOX9 (**i**) and 7 days for Aggrecan (**ii**), COL2A1 (**iii**), COL1A1 (**iv**) and COL10A1 (**v**), n=3. For all figures: data are represented as mean ± standard deviation and differences are considered significant for $p \leq 0.05$ using t-tests for pairwise comparisons (* $p \leq 0.05$, ** $p \leq 0.01$); all hydrogels functionalised with 2 mM RGD peptide to allow cell adhesion. All qPCR data was made relative to control sample with stronger elastic character (0.1 loss tangent) and normalised to GAPDH. All hydrogels were crosslinked using 100% VPM peptide.

.
